# Wildlife depredation stabilizes ecological-economic systems and can lead to improved economic outcomes

**DOI:** 10.64898/2026.09.21.753200

**Authors:** Katrina J. Davis, Roberto Salguero-Gómez

## Abstract

1. Human–wildlife conflicts over shared resources are often interpreted as direct competition, whereby resources consumed by wildlife are unavailable to people. Yet, individuals of wild populations can differ in their propensity to exploit resources, and the demographic, ecological, and economic consequences of such behavioural heterogeneity remain poorly understood.
2. We developed an integrated demographic–bioeconomic model of Atlantic cod (*Gadus morhua*), grey seals (*Halichoerus grypus*), and a coastal gillnet fishery. Stage-structured population models incorporated density dependence, predator–prey interactions, adaptive harvesting, and transitions of adult seals into a bold, depredating state. We compared four scenarios of increasing complexity over 50 years across 100 coupled demographic realisations and examined sensitivity to the energetic contribution of depredated catch, harvest levels, and social amplification of depredation behaviour.
3. Cod and seals persisted in all demographic realisations, supporting our ecological-persistence hypothesis. Contrary to our predictions, however, adding bold seals neither reduced long-term fishery performance nor destabilised the system. Median net present value was approximately three-fold greater in the full system than without depredation, and was greater in 76% of paired realisations. The full system was also the most temporally stable: median cod variability declined by 74% relative to the otherwise equivalent fishery without depredation, while seal variability declined by 45%. Behavioural depredation therefore altered predator population structure and imposed direct catch losses, but these effects were counterbalanced by indirect demographic feedbacks.
4. *Synthesis and applications.* Depredating predators should not necessarily be viewed simply as competitors that reduce contested resources. Behavioural heterogeneity can introduce compensatory pathways that alter population stability and long-term economic outcomes. Management should therefore consider not only predator abundance and immediate catch losses, but also how harvest creates opportunities for behavioural change and how those changes propagate through coupled ecological– economic systems.

## Introduction

As human activities continue to expand and intensify worldwide, we are increasingly forced to share natural resources with wild animal populations (Dirzo *et al*. 2014; Venter *et al*. 2016). Nowhere is this tension more apparent than in marine ecosystems, where commercial fisheries and marine predators often target the same diminishing fish stocks (Jackson *et al*. 2024; Tixier *et al*. 2021). While fisheries often aim for sustainable harvests to support livelihoods and economies, predators consume fish as part of their natural ecology. This overlap has created persistent conflict, especially in cases where predator populations have rebounded after decades of protection (Davis *et al*. 2021; Roman *et al*. 2015).

Marine mammals, in particular, have made striking recoveries in many parts of the world following historical declines due to hunting and exploitation (Lotze *et al*. 2006). Examples include the recovery of humpback whale (*Megaptera novaeangliae*) populations in the North Atlantic and Pacific following the cessation of commercial whaling (Bettridge 2015; Roman *et al*. 2014), and the rapid rebound of grey seal (*Halichoerus grypus*) populations in the Northeast Atlantic after protections were introduced in the mid-20th century (Vincent 2017). Together, this resurgence has led to renewed interactions with fisheries, often in the form of depredation—the removal or damage of fish by predators after those fish have already been captured by humans (Read 2008). Such interactions challenge the profitability of fisheries (Davis *et al*. 2021) and the conservation of recovering predators (Davis 2022). Pinnipeds, such as seals and sea lions, are especially implicated in this conflict due to their growing populations and frequent presence in productive coastal waters (Magera *et al*. 2013).

Depredation, broadly defined, occurs when animals appropriate resources that humans perceive as their own (Read 2008). In marine fisheries, depredation involves fish being taken from nets, lines, or traps. Analogous behaviours are well documented in terrestrial settings: raiding of crops by elephants (Hoare 1999), predation of small livestock by baboons (Kifle 2021), and large carnivores preying on larger livestock (Treves *et al*. 2016). Across these contexts, individuals that repeatedly engage in depredation are often described as ‘bold’ (Sloan Wilson *et al*. 1994)—animals more willing to take risks in human-dominated landscapes. In marine systems, depredation by bold individuals spans a diverse set of species, from seabirds (González-Solís *et al*. 2000) to cetaceans (Earl *et al*. 2021), yet pinnipeds are the most persistent actors (Graham *et al*. 2011b; Magera *et al*. 2013).

Despite the prevalence of depredation in marine environments (Tixier *et al*. 2021), we know relatively little about how it emerges and spreads–particularly in pinnipeds. Some of the best documented accounts of this behaviour are from studies of killer whales (*Orcinus orca*) depredating Patagonian toothfish (*Dissostichus eleginoides*) from long lines in the Crozet Islands (Earl *et al*. 2021). However, equivalent empirical data for pinnipeds are scarce (but see (Jackson *et al*. 2024)). While it is clear that some individuals depredate and others do not (Graham *et al*. 2011b), the mechanisms driving these behavioural transitions and their demographic impacts remain poorly understood in pinnipeds. Likewise, the extent and consequences of depredation for stable predator-prey dynamics are difficult to quantify without long-term data that explicitly capture the feedbacks between predator behaviour, prey availability, and fishery activity (Read 2008).

This knowledge gap is particularly problematic because pinnipeds are well positioned to interact frequently and intensively with fisheries. Compared to slower-growing marine predators (*e.g.*, killer whales *Orcinus orca*, albatrosses *Diomedea* spp.), pinnipeds reproduce quickly (4-6 years; (Boyd 2000a)) and many populations have rebounded rapidly from past exploitation (Davis *et al*. 2021; Magera *et al*. 2013). Their coastal breeding and haul-out behaviours further increase the likelihood of overlap with small-scale fisheries, which are more vulnerable to depredation because of the smaller scale of their activities compared to larger commercial fleets (Jackson *et al*. 2024). Recovering pinniped populations have also prompted concerns that these species contribute to the poor recovery or continued degradation of commercial fish stocks, including Atlantic salmon fisheries in Scotland (Butler *et al*. 2011) and commercial stocks in Cape Cod (Gruber 2014). In most systems, we lack reliable data on pinniped population trajectories, spatial distributions, and depredation rates—especially over time.

In the absence of rich empirical datasets, modelling offers a tractable approach to understand how different fishery-seal-prey scenarios can sustain the emergence of depredation and to quantify the impact of depredation on economic, population, and community consequences (Gaillard & Yoccoz 2003; Punt & Donovan 2007). By integrating demographic and bioeconomic models, we can explore how bold behaviours may arise in predator populations and how their emergence feeds back into fishery dynamics. Demographic models allow us to track how transitions among life cycle stages (e.g., juveniles, adults, bold adults) affect the overall dynamics of the population via direct and indirect contributions (Caswell 2001), such as density dependence (Brook & Bradshaw 2006). Bioeconomic models, in turn, link these non-linear dynamics to fishing effort, profitability, and management outcomes (Holma *et al*. 2014). Although the biological and economic links between fish, predators, and fisheries have long been acknowledged (Clark 1993), these links have rarely been formalised in models that include behavioural transitions such as depredation.

Here, we develop this model for a well-defined, ecologically and economically important system: Atlantic cod (*Gadus morhua*), the grey seal (*Halichoerus grypus*), and the UK gillnet fishery. Atlantic cod is among the UK’s most heavily exploited species (Fernandes & Cook 2013) and is an important component of grey seals’ diet (Hammond *et al*. 1994). Conflict between gillnet fisheries and seals is well documented, including in the Baltic Sea (Kindt-Larsen *et al*. 2023), where it has been estimated that 4.1 cod are lost for every cod landed. Grey seals are demographically well studied, and existing data provide estimates of their population structure, growth, and overlap with fishing activity (Bull *et al*. 2021; Tanner & Davis 2026). We model a three-actor system comprising cod stocks, a gillnet fishery, and grey seals, some of which can transition to a bold, depredating state. Our approach incorporates intra- and inter-specific density dependence in survival, growth, and reproduction for both cod and seals, while explicitly linking the emergence and demographic consequences of depredating behaviour to fishery activity.

To disentangle the effects of predation, fishing, and depredation, we simulate four scenarios of increasing complexity: (1) fish only; (2) fish and seals; (3) fish, seals, and the fishery; and (4) all actors, including bold depredating seals. Specifically, we hypothesise that (<u>H1: Ecological persistence hypothesis</u>) interspecific density-dependent and predator–prey feedbacks will promote the long-term persistence of cod and seals rather than system collapse. We further predict that (<u>H2: Non-linear fishery profitability hypothesis</u>) fishery profitability will decline non-linearly as catch increases because increasing depredation opportunities progressively increases the number of adult seals becoming bold and this reduces the proportion of harvested fish that can be landed and sold. Finally, we hypothesise that (<u>H3:</u> <u>System stability hypothesis</u>) the emergence of bold, depredating seals will destabilise the dynamics of the fish–seal–fishery system, increasing temporal variability relative to otherwise equivalent systems without depredation. Together, these hypotheses allow us to examine how behavioural heterogeneity among predators can alter not only the economic consequences of sharing a resource with wildlife, but also the dynamics and stability of the coupled ecological– economic system. By identifying the feedbacks through which fishing activity, predator behaviour, and population dynamics interact, our framework provides a basis for evaluating management strategies that seek to reduce human–wildlife conflict while maintaining both ecological and economic sustainability.

## Methods

To test our three hypotheses regarding the demographic and economic consequences of seal depredation on fisheries, we developed a multi-species, stage-structured modelling framework that dynamically integrates fish and seal population processes with fishing activity, economic returns, and behavioural transitions. The model tracks three interacting components over a 50-year time horizon: Atlantic cod (*Gadus morhua*), grey seals (*Halichoerus grypus*), and a gillnet fishery. Population dynamics for fish and seals are represented using stage-structured matrix population models (Lefkovitch 1965). However, unlike in standard formulations on population ecology (Salguero-Gomez *et al*. 2016), projection matrices are updated at each time step as functions of population state and harvesting intensity. Specifically, to test our hypotheses H1, H2, and H3, the vital rates of survival, maturation, and reproduction are modified by density-dependent and interaction-dependent functions that capture intra- and inter-specific feedbacks, as well as fishing pressure. In addition, adult seals can transition to a bold, depredating behavioural state through a state-dependent function linked to fishery catch. This feature creates feedbacks between predator behaviour and harvesting outcomes. Annual dynamics are simulated iteratively, generating time-series of population abundances, catch, depredation rates, and economic returns. All analyses were conducted in R (v. 4.2.2). Demographic data processing, matrix manipulation, phylogenetic analyses, and population projections used the R packages Rcompadre and Rage (Jones *et al*. 2022), popbio (Stubben & Milligan 2007), popdemo (Stott *et al*. 2012), rfishbase (Boettiger *et al*. 2012), Rphylopars (Goolsby & Bruggeman n.d.), phytools (Revell 2012), rotl (Michonneau *et al*. 2016), ape (Paradis *et al*. 2004),

### Scenario design and initial conditions

To examine how ecological interactions and economic feedbacks shape system dynamics, we simulated four scenarios of increasing complexity, each adding a key process to the system. In Scenario I (fish only), Atlantic cod populations were modelled in the absence of seals and fishing. Scenario II (fish + seals) introduced predator-prey interactions between cod and grey seals, but excluded fishing and depredation. Scenario III (fish + seals + fishery) incorporated harvesting by the fishery alongside seal predation of cod, but did not allow seals to depredate fisheries catch. Scenario IV (full system) included all components, allowing seals to also transition to a bold behavioural state and depredate fishery catches, thereby generating feedbacks between fishing activity and predator behaviour. This step-wise approach allows us to isolate the effects of predation, harvesting, and behavioural depredation on system dynamics. See Table 1 for full details of scenario parameterisation.

**Table 1.** We simulated four different scenarios to test three hypotheses regarding the ecological persistence (H1), non-linear fishery profitability (H2), and overall system stability (H3) of a cod-seal-fishery system. Each scenario increases complexity and realism. The table details the parameterisation of our models for each scenario. ***N****_fish_* is the population vector of the examined cod population, composed of juveniles and adults. Similarly, ***N****_seal_* is the population vector for seals, with juvenile, (non-bold) adult, and bold adult seals. *N_fishery_* is a scalar that represents the numbers of vessels in our simulations. H1 compares and contrasts outcomes of scenarios I, II, and III; H2 focuses on III and IV; and H3 on III *vs.* IV.

|  |  | Scenario |  |  |  |
| --- | --- | --- | --- | --- | --- |
| Initial conditions | Break-down | I: Only fish | II: Fish and (not bold) seals | III: Fish, (not bold) seals, and fisheries | IV: Fish, bold seals, and fisheries |
| $N_{fish}$ | Total | 15,000 | 15,000 | 15,000 | 15,000 |
|  | Juvenile | 10,000 | 10,000 | 10,000 | 10,000 |
|  | Adult | 5,000 | 5,000 | 5,000 | 5,000 |
| $N_{seal}$ | Total | 0 | 70 | 70 | 70 |
|  | Juvenile | 0 | 20 | 20 | 20 |
|  | Adult | 0 | 50 | 50 | 48 |
|  | Bold adult | 0 | 0 | 0 | 2 |
| $N_{fishery}$ | Total | 0 | 0 | 1 | 1 |

Briefly, to test our three hypotheses, we compared the temporal dynamics and long-term outcomes generated by the four scenarios, focusing on contrasts that isolate each successive ecological, economic, and behavioural process. To test the ecological persistence hypothesis (H1), we compared cod and seal population trajectories across scenarios I, II, and III to determine whether populations persist over the 50-year simulations as predator–prey-fishery interactions are introduced. To do so, we examined both total abundance and stage-specific dynamics for evidence of coexistence *vs.* collapse. To test the non-linear fishery profitability hypothesis (H2), we focused on the fishery scenarios III and IV, comparing landed catch, revenues, costs, annual profits, and long-term economic performance, and examine how these outcomes change with the abundance of bold seals and the resulting level of depredation. To test the system stability hypothesis (H3), we contrasted otherwise comparable systems without and with depredation (scenarios III and IV, respectively), assessing whether the introduction and subsequent spread of bold seals increases the magnitude and temporal variability of fluctuations in cod, seals, catch, and economic returns. Finally, we used sensitivity analyses that vary depredation rates and key interaction parameters to evaluate whether these demographic, economic, and stability outcomes are robust to alternative assumptions and to identify the conditions under which the predicted feedbacks are strengthened or weakened. Their implementation is described under “Sensitivity analyses” (below).

We assessed population persistence (H1) by recording the final and minimum total abundance reached by each population over the 50-year projection. We defined numerical extirpation as total population abundance falling below 1 individual at any point during the projection. This threshold represents an effectively absent population in the demographic model and is used as an operational diagnostic of population collapse rather than as a biologically defined minimum viable population (MVP). Because initial population sizes were selected to examine comparative system dynamics rather than to reproduce a specific empirical population (*e.g.*, (Davis 2022)), we did not impose an externally derived MVP threshold.

To enable direct comparison across the four scenarios, we initialised all simulations with identical starting conditions. Specifically, we set the cod population to a total abundance of ***N****_fish_* = 15,000 individuals, comprising 10,000 juveniles and 5,000 adults. This juvenile-skewed structure reflects typical life histories of marine fishes, where early life stages dominate population abundance due to high reproduction and mortality (*e.g.*, (Anderson *et al*. 2008; Beverton & Holt 1957)). For scenarios including seals (scenarios II–IV), seal populations were initialised at ***N****_seal_* = 70 individuals, with 20 juveniles and 50 adults, representing a small but reproductively active population consistent with the slower life histories and lower abundances of marine mammals (Boyd 2000b). In scenario IV, we initialised the bold behavioural state with 2 adult seals to allow the emergence and spread of depredation within the population. This low initial frequency reflects the assumption that depredation is initially rare and arises from a subset of individuals (Graham *et al*. 2011a). The fishery, included in scenarios III and IV, was represented by a single vessel as a simplified unit of fishing effort, thus allowing effort and economic outcomes to scale with system dynamics rather than fleet size. We chose these initial abundances to represent moderate rather than extreme population sizes, thus avoiding immediate extinction or unrealistically rapid growth, and allowing transient dynamics and feedbacks to emerge. While these values are not intended to reproduce a specific empirical system, they provide a biologically plausible starting point for exploring system behaviour. Importantly, our focus is on comparative dynamics across scenarios rather than absolute population sizes, and results were qualitatively robust to alternative initial conditions (see Sensitivity Analyses in the Supplementary Materials).

### Demographic models for fish and seals

We modelled the population dynamics of both Atlantic cod and grey seals using stage-structured matrix population models (MPMs hereafter; (Caswell 2001)) that incorporate survival (*σ*), maturation (*γ*), and reproduction (Ø) as core demographic processes. The fish (*f*) model consists of two stages: juveniles (*j*) and adults (*a*) (Fig. 1B); while the seal (*s*) model includes three stages: juveniles (*j*), adults (*a*), and bold adults (*b*) (Fig. 1C). We represented the life cycle of each species with a Lefkovitch matrix model (Lefkovitch 1965) that was dynamically updated at each time step to reflect internal (density-dependent) and external feedbacks (*e.g*., harvest and depredation).

**Figure 1.**
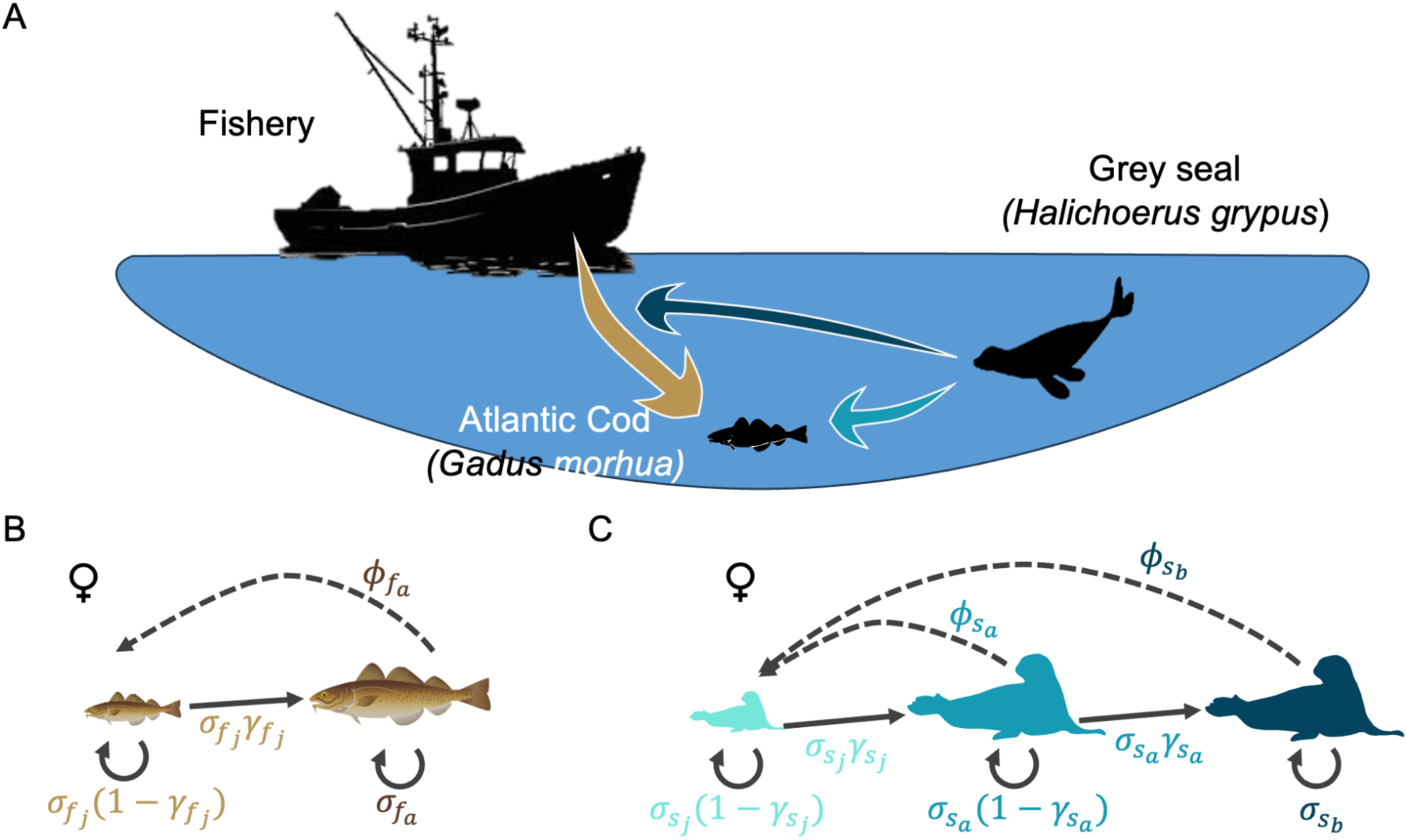
Conceptual and demographic structure of our modelling framework to examine the effects of depredation in a fishery system. **A**. Schematic representation of interactions between Atlantic cod (*Gadus morhua*), grey seals (*Halichoerus grypus*), and a coastal gillnet fishery. Seals can predate cod directly (light green arrow) or by consuming part of the fishery catch (yellow arrow) via depredation (dark green arrow). **B**. Two-stage, female-only life cycle of the fish (*f*) Atlantic cod, comprising juveniles (*j*) and adults (*a*). **C**. Three-stage, female-only life cycle of grey seals (*s*), comprising juveniles (*j*), adults (*a*), and bold adult individuals (*b*). Bold adults represent individuals that engage in depredation of fishery catch. In both cod and seal: *σ* = survival; *γ* = maturation; and Ø = reproduction.

#### Atlantic cod

To simulate the dynamics of fish-seals-fishery interactions, we identified the Atlantic cod (*Gadus morhua*) as an ideal target fish species for both seals and fisheries. Atlantic cod is of high commercial importance in many countries (Worm *et al*. 2009), and has experienced severe declines in recent decades (Hutchings & Myers 1994). Importantly, Atlantic cod has been recorded as having been depredated by grey seals (Glemarec *et al*. 2024; Tanner & Davis 2026). A key area where this species is commercially targeted is the North Sea, where populations declined 70% between 1985-2003, and are now considered vulnerable to extinction (Crilly & Esteban 2013).

High-resolution, stage-structured demographic information for Atlantic cod does not exist. Thus, we modelled its dynamics based on phylogenetic imputations of closely related species and for a relatively simple model with only two stages: juvenile and adult cod (Fig. 1B). This model includes four vital rates: juvenile fish survival (*σ_fis_*_ℎj_), fish maturation (*i.e.,* survival-independent transition probability to adulthood, *γ_fis_*_ℎj_), adult survival (*σ_fis_*_ℎ_*_a_*), and adult reproduction (*φ_fis_*_ℎ_), as shown in the MPM ***A****_fish_* in Equation 1.

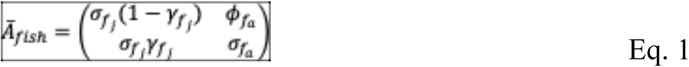

The vital rates of survival, maturation, and reproduction are typically under strong selection, and so they often have a high phylogenetic signal (Blomberg *et al*. 2003; James *et al*. 2021)). This is a key condition for robust phylogenetic imputations (Blomberg *et al*. 2003; Freckleton & Orme 2002; Penone *et al*. 2014). To enable phylogenetically informed imputation of vital rates for cod (*G. morhua*), we first assembled a comparative dataset of demographic models for related species. To that end, we obtained MPMs for ray-finned fishes (Class Actinopterygii) from the COMADRE Animal Matrix Database v. 4.21.1 (Salguero-Gómez et al. 2016). Using the R package Rcompadre (Jones *et al*. 2022), we imposed a series of criteria to retain vital rate data fit for subsequent analytical steps. First, we only retained MPMs for studies where the examined fish species was studied in the wild and under unmanipulated conditions to represent natural population dynamics. We further excluded MPMs lacking reproductive information (submatrix ***F****_fish_* in Eq. 2), or for which the survival-dependent components (submatrix ***U****_fish_* in Eq. 2) and reproductive-dependent components could not be separated explicitly. Finally, to reduce sampling bias, we retained only one MPM per species. The result set contained MPMs for 65 fish species (Table S1, Figure S1). Here, the MPM ***A*** summarised the population dynamics across the full length of the study.

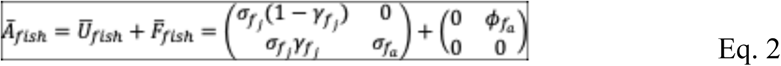

To facilitate the imputation of the four vital rates involved in our Atlantic cod two-stage MPM (Eqs. 1,2), we modified the set of MPMs from the resulting 65 species from the previous steps. The matrix dimensions (*i.e.*, the number of stages in the life cycle stage of the species model) of these species ranged from 2 to 11 (Figure S2). Accordingly, we next standardised these MPMs to a two-stage juvenile–adult structure (as in Eq. 1) using methods described in Salguero-Gómez & Plotkin (2011) via the function mpm_collapse from the R package Rage (Jones *et al*. 2022). Briefly, the collapsing algorithm preserves the eigenstructure of the original MPM, while allowing for the collapse of an MPM to any dimension. Following Salguero-Gómez and Plotkin (2010), we collapsed the original MPM leaving the first class unaltered, and collapsing all others, as this approach retains the original eigenstructure of the source MPM.

To provide more phylogenetic strength to our imputations in the next step, we obtained adult body mass information. The rationale is that adult body mass is an excellent correlate of vital rates (James *et al*. 2021) and life history traits (Gaillard *et al*. 1989; Healy *et al*. 2019). We obtained adult body mass values from FishBase (Froese & Pauly 2002) via the R package rfishbase (Boettiger *et al*. 2012). We subsequently log-transformed body mass to adhere to normality assumptions in the next analytical steps. Finally, we added to the result dataset our target species Atlantic cod (*Gadus morhua*), with adult body mass also obtained from FishBase, but naturally with the missing values for the four vital rates shown in Eq. 1.

To provide a phylogenetic framework for imputation, we constructed a species-level phylogeny for our 66 fish species: 65 from COMADRE plus *Gadus morhua*. As a pre-step, we checked for the need to update the scientific names of the species using taxize (Chamberlain & Szocs 2013) and then matched them using the Open Tree of Life taxonomy via rotl (Michonneau et al. 2016). We assigned the branch lengths of the resulting phylogenetic tree using the function compute.brlen from the R package ape (Paradis et al. 2004). The resulting tree was confirmed to be ultrametric and pruned to match our dataset. To provide reassurance of the phylogenetic imputation of the vital rates of Atlantic cod, we first assessed the phylogenetic signal for body mass and the four examined vital rates using Blomberg’s *K* (Blomberg *et al*. 2003) and Pagel’s *λ* (Pagel 1999). To do so, we used the function phylosig from the R package phytools (Revell 2012). All traits showed strong phylogenetic signals (Pagel’s *λ*: 0.803-0.927; Blomberg’s *K* > 0.5; Table S2), supporting the assumption of phylogenetic conservatism. These estimates justified the use of phylogenetic information to estimate missing trait values (Penone *et al*. 2014; Revell *et al*. 2008).

To impute the missing vital rates of *Gadus morhua*, we used multivariate phylogenetic imputation implemented with the phylopars function in the R package Rphylopars (Goolsby & Bruggeman n.d.). The analysis incorporated juvenile survival, juvenile maturation, adult survival, adult reproduction, and adult body mass (Table S1), with the survival and reproduction traits log-transformed prior to imputation and subsequently back-transformed, under the assumption of Brownian motion. To propagate uncertainty associated with the composition of the comparative dataset, we generated 40 bootstrap datasets by resampling 63 fish species with replacement while retaining *G. morhua* as the focal species in each dataset. For each bootstrap replicate, the phylogeny was pruned to the species included in that replicate and phylopars was used to estimate the missing trait values for *G. morhua*. We summarised the 40 imputed values for each vital rate using their mean and empirical 95% interval (Table S3), and then generated 100 parameter realisations by independently drawing each vital rate from a uniform distribution bounded by its corresponding 95% interval. Each realisation was used to construct a two-stage, 2 × 2 matrix population model for *G. morhua* (as per Eqs. 1-2), yielding 100 alternative cod MPMs that were carried forward into the subsequent demographic simulations.

We validated the resulting Atlantic cod MPM by calculating key demographic metrics for the bootstrapped 100 MPMs. These demographic metrics include population growth rate (*λ*), generation time (*T*), net reproductive output (*Ro*), and age at maturity (*La*). We estimated these metrics using functions from popdemo (Stott et al. 2012) and Rage (Jones *et al*. 2022). The resulting model provided a plausible and internally consistent approximation of *G. morhua* demography (Fig. S3). Specifically, the estimates of population growth rate (*λ* = 1.507 ± 0.417 S.D.), mean life expectancy (*L_e_* = 6.570 ± 3.891 years), age at maturity (*L_a_* = 2.006 ± 0.005 years), generation time (*T* = 4.050 ± 1.611), and net reproductive output (*R_o_* = 5.682 ± 5.414), all fall within the range of values reported for this species (Hutchings & Myers 1994; Olsen *et al*. 2005).

To explicitly accommodate intra-specific competition and predatory effects of seals on the fish vital rates, each vital rate was modified at each time step as a function of the values of the cod and seal populations. See “Temporal dynamics, density-dependence and behavioural transitions” below.

#### Grey seal

To simulate the dynamics of a seal population, we chose the grey seal (*Halichoerus grypus*). The UK is home to the largest population of grey seals in the world (Bowen & Lidgard 2013) and existing research describes grey seal depredation rates of fisheries catch (Tanner & Davis 2026), which include Atlantic cod. We used the three demographic studies available for this species in COMADRE v. 4.21.1 (Bull *et al*. 2021; Harwood & Prime 1978; Holma *et al*. 2014). Each empirical MPM was collapsed to two biologically meaningful stages, juveniles and adults, using the matrix-collapsing approach described above. From each resulting 2 × 2 matrix population model, we extracted juvenile survival, juvenile maturation, adult survival, and adult reproduction using the same definitions as for the cod MPM. These three empirical estimates were then used to characterise uncertainty in the baseline seal vital rates. Because survival and maturation are bounded between 0 and 1, we fitted beta distributions to the observed values using method-of-moments estimates; adult reproduction, which is positive and unbounded, was represented by a lognormal distribution parameterised from its empirical mean and standard deviation. We drew 100 independent combinations of the four vital rates from these distributions and constructed a corresponding two-stage MPM for each realisation.

To capture the emergence of depredation behaviour, we extended this MPM by further partitioning the adult class into two distinct behavioural states: non-bold adults and bold adults (Figure 1C). Bold individuals are defined as those that engage in depredation of fishery catches (Read 2008), a behaviour widely documented across marine predators exploiting fishing gear (*e.g*., pinnipeds, cetaceans, and seabirds; (González-Solís *et al*. 2000; Magera *et al*. 2013; Tixier *et al*. 2015)). This extension results in a three-stage model comprising juveniles (*j*), non-bold adults (*a*), and bold adults (*b*), where we explicitly model the vital rates of seal survival (*σ_seal_*), maturation (*γ_seal_*), and reproduction (*φ_seal_*) as described in equations 3 and 4.

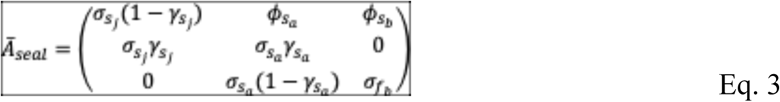

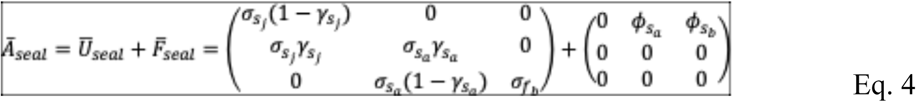

The inclusion of a bold stage is motivated by both empirical and theoretical considerations. First, depredation is typically performed by a subset of individuals within populations rather than uniformly across all adults, indicating consistent among-individual behavioural differences (Earl *et al*. 2021; Graham *et al*. 2011b). Such heterogeneity is well described within the broader framework of animal personality and behavioural syndromes, where “bold” individuals are more likely to exploit risky, human-associated resources (Sih *et al*. 2004; Sloan Wilson *et al*. 1994). Second, depredation behaviour can be learned and socially transmitted, leading to its spread within populations over time (Earl *et al*. 2021; Tixier *et al*. 2015). Representing this behaviour as a discrete stage therefore allows us to capture both its demographic consequences and its potential to increase in frequency through state transitions.

We modelled the transition between non-bold and bold adult states as a catch-dependent probability (*γ_seala_*), such that exposure to fishery harvest in the previous time step increases the probability that a surviving non-bold adult enters the bold state in the current time step. Specifically, this transition probability was modelled as an increasing saturating function of previous-year harvest, H_t-1_, with a parameter *β* controlling the sensitivity of behavioural transition to fishery activity, thus reflecting how depredation emerges when anthropogenic foraging opportunities become sufficiently profitable for the animal (Read 2008; Tixier *et al*. 2015). This assumption is consistent with optimal foraging theory, whereby individuals adopt strategies that maximise energetic returns relative to effort and risk (Stephens & Krebs 2019). Indeed, increased fishing effort or harvest effectively lowers the cost of prey acquisition by concentrating and immobilising fish, thereby incentivising seals to switch from foraging to depredation (Westphal *et al*. 2025). Conversely, the probability of remaining in the non-bold adult state decreases as harvest increases, reinforcing the behavioural feedback between fisheries activity and predator strategy. This formulation captures the empirical observation that depredation rates tend to increase with fishing intensity and repeated exposure to fishing gear (Jackson *et al*. 2024; Tixier *et al*. 2015). We further assume that bold adult individuals do not retrogress to non-bold adult individuals regardless of fishery conditions (but see Discussion).

Once individuals transition into the bold adult state, we allow their vital rates to differ from those of non-bold adults to reflect the energetic and demographic consequences of depredation. To do so, we assumed that bold adults experience a slight decrease (×0.98) in survival (*σ_sealb_*), but a slight increase (×1.01) in reproduction (*φ_seala_*) under high catch conditions compared to the survival (*σ_seala_*) and reproduction (*φ_seala_*) of non-bold adults. This key difference represents a starting point in the model regarding the predictable risks *vs.* energy intake obtained from depredating fish already captured in gear. Indeed, similar demographic benefits of anthropogenic food subsidies have been documented across taxa, including marine mammals, seabirds, and terrestrial carnivores (Newsome *et al*. 2015; Oro *et al*. 2013). For pinnipeds specifically, access to concentrated prey resources has been linked to improved body condition and reproductive output (Bowen & Lidgard 2013; McConnell *et al*. 1992). Analogous patterns have also been observed in depredating killer whales, where individuals exploiting fisheries exhibit altered foraging strategies and potentially enhanced energetic gains (Earl *et al*. 2021; Tixier *et al*. 2015). The underlying vital rates and similar emerging population growth rates of the seal model without *vs.* with bolds are shown in Figure S4.

Importantly, we assume that bold and non-bold adults share the same baseline demographic structure (*i.e*., maturation pathways and underlying life history constraints), and differ only in the magnitude of their vital rates as modified by fishery-dependent functions (Table 2). This assumption ensures that behavioural differences influence population dynamics through realistic demographic pathways rather than imposing structurally distinct life cycles.

**Table 2.** Parameterisation of density-, interaction-, and harvest-dependent vital rates in the Atlantic cod (*Gadus morhua*)–grey seal (*Halichoerus grypus*) model. *v_0_* denotes the baseline vital rate. *N_fis_*_ℎj_ and *N_fis_*_ℎ_*_a_* are juvenile and adult cod abundances; *N_seal_* _j_,*N_seal a_* and *N_seal b_*are juvenile, non-bold adult, and bold adult seal abundances, respectively; *H* is total fishery catch and *D* is catch removed through depredation; *g_Neg_*, *g_Pos_*, *g_D_*, and *g_J_* are the abundance-dependent response functions defined in Eqs. 19–22, and *h* is the catch-dependent response defined in Eq. 23. *K* controls the abundance scale of the response and *α* its magnitude. Unless otherwise specified, drivers are evaluated at the previous annual time step, *t-1*. Vital rates are: survival (*σ*), maturation (*γ*), and reproduction (*φ*).

| Species | Life cycle stage | Vital rate | Reference equation | $\alpha / \alpha_H$ | $K / \text{density scale}$ | $\beta_H$ | Min–max | Stochastic SD |
| --- | --- | --- | --- | --- | --- | --- | --- | --- |
| Fish | Juvenile | Survival $\sigma$ | Eq. 24 | 0.800; 0.900 | 5000; 10 | — | 0.010–0.400 | 0.0005 |
| | Adult | Survival $\sigma$ | Eq. 25 | 0.590; 0.950 | 5000; 10 | — | 0.300–0.970 | 0.0005 |
| | Juvenile | Maturation $\gamma$ | Eq. 26 | 0.015; 0.050 | 3000; 20 | — | 0.500–0.960 | 0.0005 |
| | Adult | Reproduction $\phi$ | Eq. 27 | 0.005; 0.010 | 7500; 20 | — | 5–60 | 0.0010 |
| Seal | Juvenile | Survival $\sigma$ | Eq. 28 | 0.150; 0.300 | 5000; 10 | — | 0.010–0.970 | 0.0500 |
| | Non-bold adult | Survival $\sigma$ | Eq. 29 | 0.050; 0.950 | 5000; 10 | — | 0.100–0.990 | 0.0500 |
| | Bold adult | Survival $\sigma$ | Eq. 30 | 0.005; 0.010 | — ; 10 | 0.010 | 0.010–0.991 | 0.0300 |
| | Juvenile | Maturation $\gamma$ | Eq. 31 | 0.040; 0.120 | 5000; 20 | — | 0.010–0.990 | 0.0005 |
| | Non-bold adult | Transition $\gamma$ | Eq. 32 | 0.035; 0.080 | — ; 10 | 0.015 | 0.000–0.990 | 0.0015 |
| | Non-bold adult | Reproduction $\phi$ | Eq. 33 | 0.080; 0.350 | 3000; 10 | — | 0–1 | 0.0500 |
| | Bold adult | Reproduction $\phi$ | Eq. 34 | 0.100; 0.100 | — ; 10 | 0.001 | 0–1 | 0.0500 |

### Fishery & bioeconomic model

We modelled a simplified representation of the UK gillnet fishery (scenarios III and IV) targeting Atlantic cod in the North Sea. We modelled the fishery as a single decision-making unit whose harvesting behaviour responds dynamically to fish population growth and economic opportunities. The UK fleet accounted for approximately one third of North Sea cod landings in the early 2000s (Fernandes & Cook 2013), making it a historically important actor in this system. We focus on gillnets because passive fishing gear is susceptible to depredation by marine predators, including pinnipeds, due to the prolonged availability and accessibility of captured fish (Jackson *et al*. 2024).

Gillnets are vertically suspended in the water column using floats and weights, passively entangling fish that encounter the gear. This fishing method disproportionately targets adult cod, with modal catch ages around three years and an average mass of approximately 2 kg per individual in the North Sea (Crilly & Esteban 2013). Accordingly, fishing mortality was applied only to the adult component of the cod population (*a* in Figure 1B), while juveniles (*j*) were not directly harvested. This assumption reflects both gear selectivity and fisheries regulations that limit the capture of immature individuals. We represented fishing effort implicitly through an adaptive, surplus-production harvest rule, broadly reflecting the principle underlying maximum sustainable yield (MSY), whereby harvest is supported by renewable population production (“Quantitative fisheries stock assessment: Choice, dynamics and uncertainty” 1992; Schaefer 1991). Fishing occurred only in scenarios III and IV. At each annual time step, the cod population was first projected forward according to the demographic model, producing the adult population available immediately before fishing, *N_prefis_*_ℎ_*_a_*_,*t*_. From year 3 onwards potential harvest *H*^∗^ was then defined as the positive increase in adult abundance relative to the adult population remaining after harvest in the preceding year, *N_postfis_*_ℎ_*_a_*_,*t*–1_:

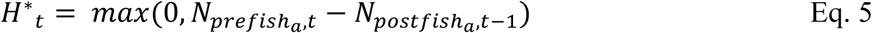

The realised harvest was additionally constrained by the number of adult cod available before fishing:

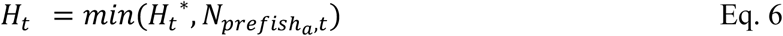

Thus, when the adult population increased between annual time steps, the fishery removed the corresponding increase in adult abundance; when adult abundance failed to increase, no harvest occurred. Importantly, harvest was calculated from the population projected before the current year’s fishing mortality was imposed, preventing previous harvest removals from being interpreted as negative population growth in the subsequent year. This formulation maintains the adult stock at approximately its preceding post-harvest abundance when population growth permits, while allowing the stock to decline without additional fishing mortality when demographic production is insufficient. It therefore represents a simplified surplus-production harvesting rule rather than an explicit implementation of MSY.

We next incorporated economic dynamics through a bioeconomic submodel linking cod abundance, fishing costs, depredation, revenue, and profit. We calculated the cost of harvesting an individual cod as a stock-dependent, bounded exponential function:

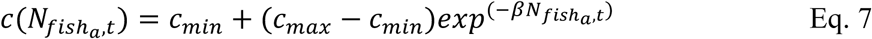

where *N_fis_*_ℎ_*_a_*_,*t*_ is adult cod stock abundance, *c_min_* and *c_max_* are the lower and upper unit-cost bounds, respectively, and *β* determines the rate at which unit harvesting costs decline with increasing stock abundance. This formulation represents increasing fishing costs per fish as cod abundance declines because lower stock abundance reduces fishing efficiency and requires greater effort to obtain a given catch.

The lower and upper unit-cost bounds were informed by empirical cost estimates for the UK <12m gillnet fleet reported by Crilly and Esteban (2013). The lower bound comprised effort-related operating costs, including fuel, crew, variable operational costs, and repair and maintenance, representing the costs associated with active fishing operations. The upper bound included all reported operating, fixed, and capital costs, representing a high unit-cost operating regime in which the costs of maintaining fishing capacity are distributed across a lower level of catch. This approach mirrors the greater effort required to locate and capture fish at low densities (Squires 1987). Costs were converted from £ tonne^-1^ to £ fish^-1^ assuming a mean cod mass of 2 kg. We parameterised *β* such that the unit cost was halfway between *c_min_* and *c_max_* when adult cod abundance reached one-half of the reference equilibrium stock, giving:

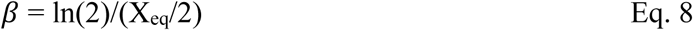

where the reference equilibrium stock, X_eq_, was calculated as the median total cod abundance at the end of the fish-only scenario I. Total fishing costs were then calculated as the unit cost at the current adult cod abundance multiplied by the total catch before depredation. This formulation provides a generic representation of the expected decline in harvesting efficiency as cod abundance decreases without requiring estimation from a time series of catch, effort and price data. Total fishing costs were then calculated as the stock-dependent cost per fish multiplied by total fish caught before depredation:

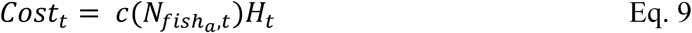

Thus fishing costs are incurred for the full harvest, including fish subsequently removed through depredation. We estimated depredation, *D_t_*, based on the annual energetic requirements of an adult female grey seal. We assumed a mean body mass of 117.4 kg, based on measurements of actively foraging adult female grey seals in eastern England (Armstrong *et al*. 2023). Daily energetic requirements were then estimated following the bioenergetic approach used by (Gardmark *et al*. 2012). Basal metabolic rate (BMR, kJ individual^-1^ day^-1^) was calculated as:

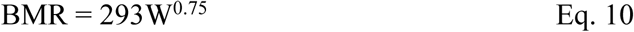

where *W* is body mass (kg). Following Gardmark et al. (2012), we assumed that energetic costs of activity were twice BMR, that 85% of consumed energy was metabolizable, and that 10% was lost through the heat increment of feeding. These authors used this approach to estimate a daily energy requirement of 25 MJ for a 100-kg grey seal. Applying the same formulation to a 117.4 kg adult female resulted in an estimated daily energetic requirement of approximately 27.9 MJ individual^-1^ day^-1^, corresponding to an annual requirement of approximately 10.2 GJ individual^-1^ yr^-1^.

Cod energetic density was parameterised using measurements of whole, 3-year-old Atlantic cod reported by Holdway and Beamish (Beamish n.d.). We used an energetic density of 4.55 MJ kg^-1^ wet mass (converted from 1,088 cal g^-1^), and harvested adult cod were represented by individuals of mean mass 2 kg. Each harvested cod therefore contained approximately 9.10 MJ. Under the simplifying assumption that cod were the sole prey source, a bold adult female grey seal would consequently require approximately 1,119 2 kg cod yr^-1^ to meet its entire annual energetic requirement. Because depredation of fisheries catch is unlikely to provide the entirety of a seal’s annual energetic intake, we introduced a parameter, *m_D_*, representing the proportion of annual energetic requirements obtained through fisheries depredation. In the baseline model, we assumed *m_D_* = 0.1, such that fisheries depredation supplied 10 % of the annual energetic requirements of a bold seal. This corresponded to approximately 112 cod, or 224 kg of cod, depredated per bold seal per year. We evaluated uncertainty in this assumption by varying *m_D_* across alternative values in sensitivity analyses. Depredation was subsequently applied to the total catch, with the number of fish removed from fishery harvest determined by the abundance of bold seals (*N_sealb_*_,*t*_):

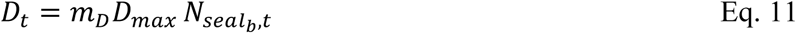

where *D_max_* is the number of 2 kg cod required to meet the full annual energetic requirements of an adult female grey seal. Realised depredation was constrained so that *D_t_* could not exceed the total fishery catch available in year *t*. The realised landings available for sale were calculated as the harvest remaining after depredation:

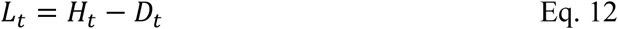

We calculated fishery revenue as:

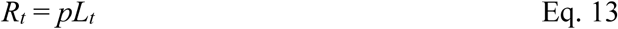

where *p* is the assumed constant market price per cod, here £2.83, and *L_t_* are the realised landings after depredation. This price assumes a 2 kg individual, and is based on prices for cod landed by the UK fleet during 2006 and 2007–the same time period as our cost data (Anderson & Guillen 2009). We assume stable cod prices in integrated European markets. Profit was calculated as:

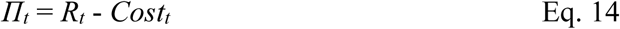

This formulation means that depredation generates a direct economic cost to the fishery because fishing costs are incurred for the full harvest, whereas revenue is received only for fish remaining after depredation. To quantify longer-term economic performance, we calculated the present value (*PV*) of annual profit in each year as:

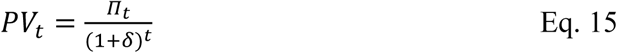

Where *δ* = 0.05 is the annual discount rate, and summed these values across the full 50-year projection to obtain total net present value (NPV). This provided a cumulative measure of fishery economic performance across the projection period.

### Temporal dynamics, density-dependence and behavioural transitions

To test our hypotheses, we projected population dynamics for both Atlantic cod and grey seals in discrete annual time steps over a time horizon of 50 years. To do so, we used the chain rule (Caswell 2001), where the population vector in the next time step (***N****_t+1_*) can be calculated as:

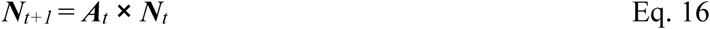

where ***A****_t_* is a time-varying projection matrix composed of survival and transition (***U***) and reproduction (***F***) submatrices (Eqs. 2 & 4), and ***N****_t_* is the stage-structured population vector at time *t* (Table 1).

Each scenario was projected for 50 annual time steps using 100 cod demographic realisations and 100 corresponding grey seal demographic realisations. The same realisation index was paired across species, yielding 100 coupled demographic realisations per scenario. We ran 400 models in total across the four scenarios. At each time step, the relevant matrix population model was updated according to current population abundance, interspecific interactions, fishery activity, and behavioural state before projecting the population to the following year.

Unlike classical MPMs with fixed vital rates (Salguero-Gomez *et al*. 2016), the vital rates in our model were dynamically updated at each time step as functions of population densities, species interactions, and fishery activity (Table 2). Indeed, a key feature of our model is that each vital rate depends not only on total population sizes, but also, when biologically pertinent, on the abundance of specific life stages within and across species. For instance, adult seals predate on both juvenile and adult fish, but with a preference for adult fish, while bold adult seals only consume adult fish, as that is the fish stage that gillnets target (Crilly & Esteban 2013). We formalise this complex relationship by defining each vital rate *v_i,t_* for stage *i* at time *t* as:

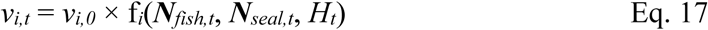

where *v_i,0_* is the baseline (density-independent) value of the vital rate, ***N****_fish,t_* and ***N****_seal,t_* are the relevant stage-specific fish and seal abundances, and *H_t_* is fishery catch. Each *f_i_*(⋅) was constructed as the product of component functions describing the relevant density-, predation-, resource-, and catch-dependent effects:

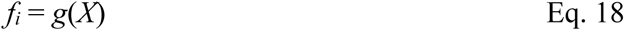

We represented abundance-dependent effects using a general saturating response function, *g*(*X*), where *X* is the abundance of the relevant fish or seal stage after rescaling by a stage- and interaction-specific density constant, *K* (*i.e*., *X*’=*X*/*K*). Depending on the biological mechanism represented, *g*(*X*) could take one of four forms:

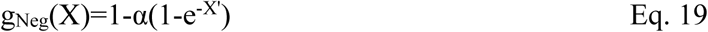

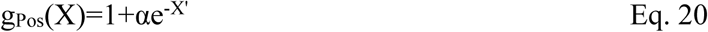

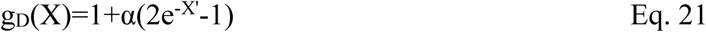

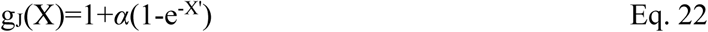

where α determines the magnitude of the response to the driver and K determines the abundance scale over which the response approaches its asymptote. The “Neg” functional form describes an effect that increasingly reduces the focal vital rate as abundance increases, but is absent when the driver is absent, whereas “J” produces the corresponding increasing response. The “Pos” and “D” forms additionally allow the focal vital rate to be elevated at low driver abundance and to decline towards its baseline (Pos) or below its baseline (D) as the driver increases. When more than one abundance-dependent driver affected a vital rate, their respective g(X) terms were multiplied before modifying the baseline vital rate. To incorporate temporal environmental stochasticity, selected vital rates were additionally multiplied at each time step by a normally distributed random deviate with mean 1 and vital-rate-specific standard deviation (S.D.), after which the resulting vital rate was constrained to its biologically specified minimum and maximum; values of stochastic S.D. are reported in Table 2.

We used a separate saturating function, *h*(*H*), for vital rates directly affected by fishery catch or depredation:

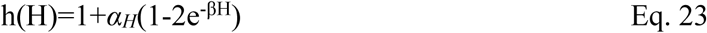

where *H* is the relevant fishery catch, *α_H_* determines the maximum magnitude of the catch-mediated effect, and *β* determines the rate at which the response approaches its asymptote. In contrast to *g*(*X*) (Eqs. 20-23), this formulation explicitly represents a shift from a negative effect when no catch is available, h(0)=1-*α_H_*, to a positive effect as catch increases, approaching 1+*α_H_* at high catch. This function was used to represent the demographic consequences of access to fishery-derived resources for bold seals. Where catch-mediated effects acted alongside density dependence, *h*(*H*) was multiplied by the appropriate seal-density response *g*(*X*) before modifying the baseline vital rate. As for *g*(*X*), resulting vital rates were constrained to biologically specified bounds.

For the fish population, the density- and harvest-dependent vital rates are:

- Juvenile fish survival is affected by adult fish abundance and by the abundance of juvenile and non-bold adult seals the previous year, with juvenile seals contributing one-third of the effective predator abundance:

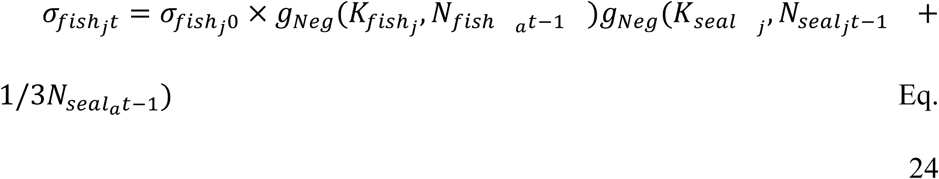

- Adult fish survival is affected by adult fish abundance and adult seal abundance:

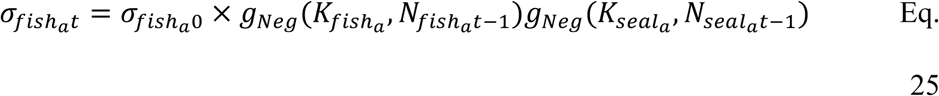

- Juvenile fish maturation to adult fish is affected by adult fish abundance, juvenile and adult seal abundance, again with the effect of adult seals being three times larger than that of juvenile seals’, and catch in the previous year, which represents the reported compensatory acceleration of cod under increased mortality pressures (Olsen *et al*. 2004):

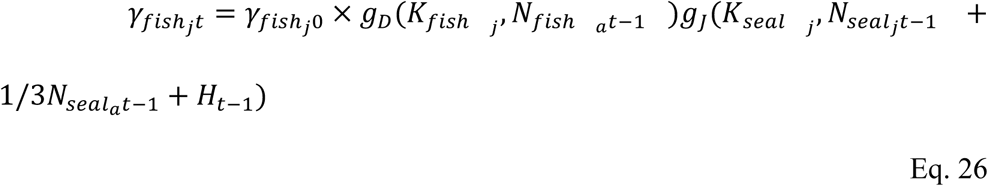

- Adult fish reproduction is affected by adult fish, adult seal abundance, and fishery catch, which represents the displacement towards a more reproductive life history when affected by increasing mortality (Kruse 2008):

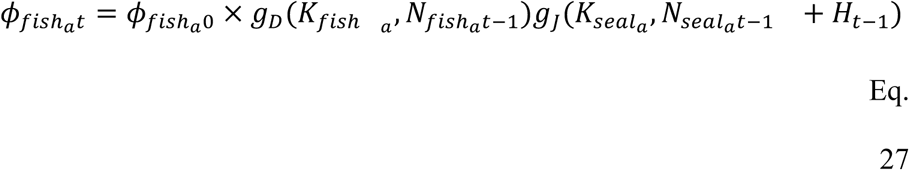

For the seal population, the density- and harvest-dependent vital rates are:

- Juvenile seal survival is affected by juvenile fish availability as well as by adult and bold adult abundances in the previous year:

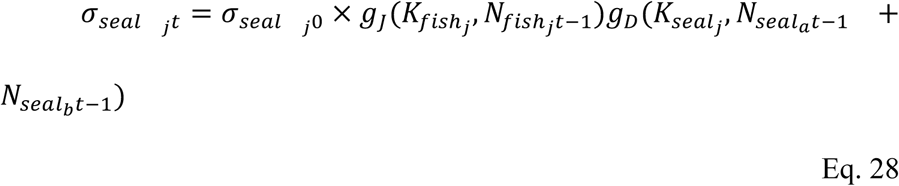

- Adult seal survival depends on fish availability, and on competition with non-bold and bold adults:

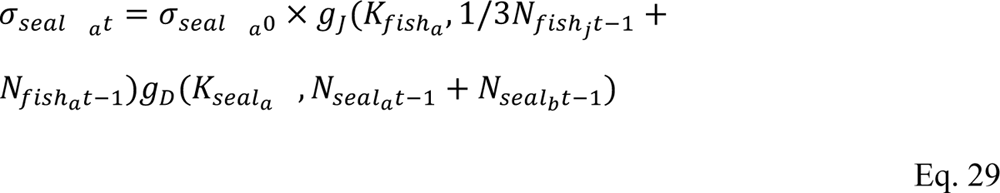

- Bold adult survival depends on the amount of bold adult seals and the (adult fish) catch depredated in the previous time step (*D_t-1_*), representing an energetic subsidy associated with access to fishery catch (Tixier *et al*. 2015):

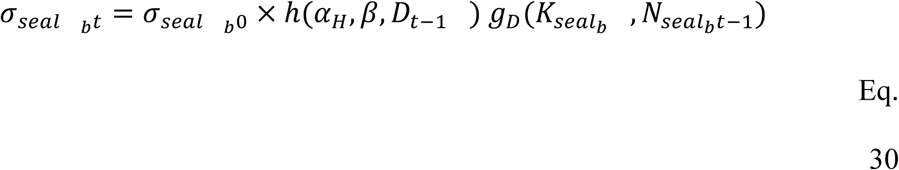

- Juvenile seal maturation depends on effective fish availability as well as adult and bold seal abundance the previous year:

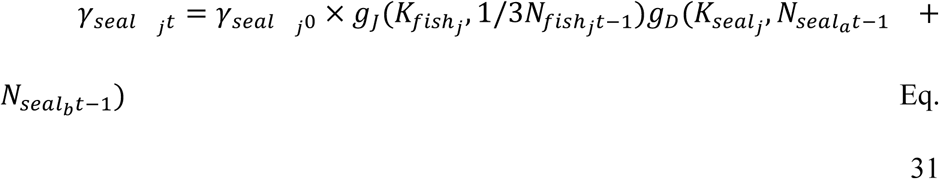

- Non-bold adult o bold adult transition depends on the total fishery catch in the previous time step, *H_t-1_*, together with the positive effect of abundance of adult bold seals, which represents both exposure to fishing and potential social or frequency-dependent amplification of depredation behaviour (Auguin *et al*. 2025):

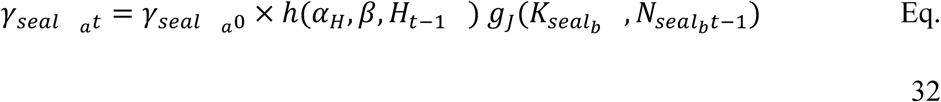

- Adult seal reproduction depends on juvenile and adult fish availability, and is impacted by adult seal competition:

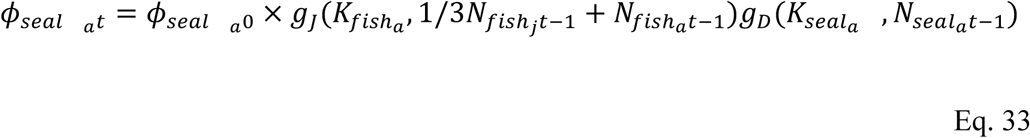

- Bold adult reproduction depends on the amount of catch depredated in the previous time step (*D_t-1_*), representing an energetic subsidy associated with access to fishery catch.

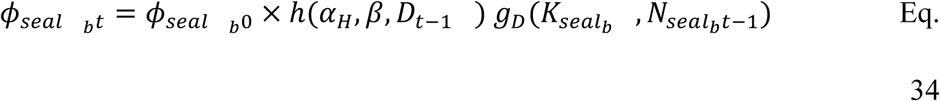

Thus, fishery activity affects the emergence of bold behaviour through exposure to total harvest, whereas the subsequent demographic performance of bold individuals depends on the amount of catch actually obtained through depredation. Together, these abundance-, behavioural-, and fishery-dependent responses create feedbacks among predator behaviour, prey dynamics, and fishery outcomes. The complete parameterisation, including response directions, sensitivity coefficients, density-scaling constants, and vital-rate bounds, is provided in Table 2.

### Sensitivity analyses

We evaluated the sensitivity of model outcomes to three assumptions that directly determine the strength of the fishery–depredating-seal feedback. First, we varied *m_D_*, the proportion of a bold seal’s annual energetic requirement assumed to be obtained through depredation, from 0.1 to 1.0 in increments of 0.1, with the baseline value of 0.1 retained as the reference. Second, we varied the adult-fish growth threshold at which fishing was initiated among 0.95, 1.00 and 1.05, representing higher, baseline and lower fishing pressure, respectively. Third, we varied the sensitivity of the transition from non-bold to bold adult seals to the abundance of existing bold adults (*sensNseal* in Eq. 33) among 0, 0.5 and 1.0, representing no social or frequency-dependent amplification, the baseline parameterisation, and a stronger amplification, respectively. Each sensitivity analysis was conducted independently while holding all other parameters at their baseline values. For each parameter value, we repeated the relevant model scenarios using the same 100 demographic realisations and stochastic random seed as the baseline analysis. We summarised effects on final fish, total seal and bold-seal abundance, cumulative catch, depredation, realised landings, cumulative net present value, and temporal variability, reporting medians and 95% simulation intervals. We did not use inferential significance tests because the model trajectories represent demographic and environmental realisations rather than independent empirical replicates.

## Results

### Hypothesis 1: Ecological persistence hypothesis

We found general support for our hypothesis H1, that interspecific density-dependent and predator–prey feedbacks would permit long-term persistence of cod and seals rather than system collapse over the 50-year projection. To test H1, we compared total and stage-specific population trajectories across scenarios I (no interspecific effects) and II (fish-seal interspecific effects; Table 1) over 50 years and quantified final and minimum abundances, together with the frequency of numerical extirpation. Neither cod nor seals underwent numerical extirpation in any of the 100 demographic realisations under any scenario. Cod abundance nevertheless differed markedly between both scenarios (Figure 2A & B). The temporal trajectories reflected key differences: cod increased to high abundance in Scenario I (only fish; Figure 2A), and showed sustained oscillations with seals in Scenario II (fish and seals; Figure 2B). Specifically, in scenario I, median cod abundance at year 50 was 4.1 million individuals (95% simulation interval: 1.25–5.84 million). Introducing seals (Scenario II) reduced median final abundance to 1.15 million (98,600–3.62 million). However, differences in abundance were not significant between the paired observations of these scenarios (*t*_99_ = 22.842, P < 0.001). In Scenario III (Figure 2C) and IV (Figure 2D), cod remained at substantially lower abundances compared to Scenarios I and II following the introduction of fishing and bold seals IV. Median final abundance was 25,859 individuals (1,324–69,813) in Scenario III. In the full system (Scenario IV), median final abundance was four-fold higher than in Scenario III, at 107,226 individuals (1,180–204,580).

**Figure 2.**
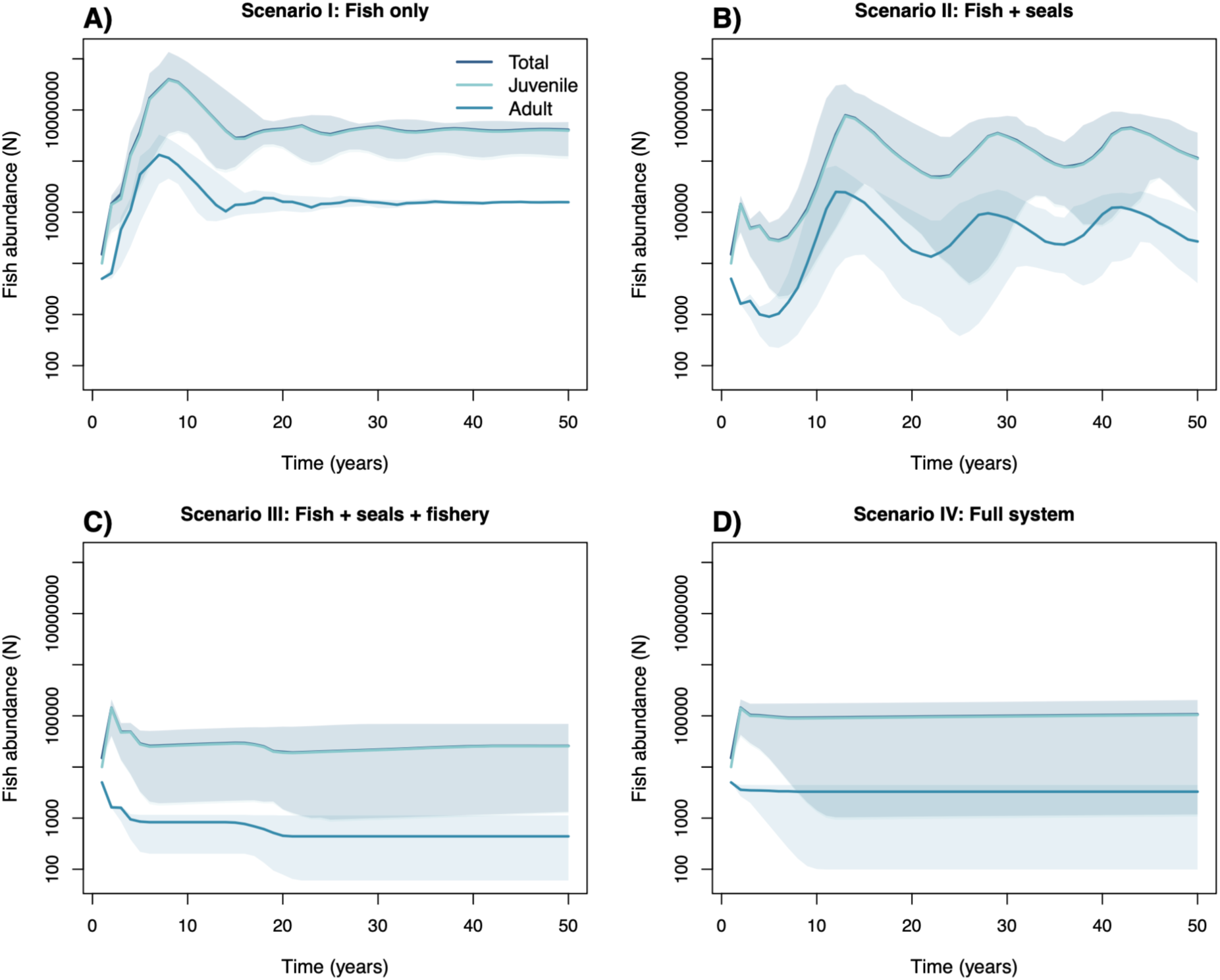
Fish population dynamics across four scenarios of increasing system complexity. Temporal dynamics of total, juvenile, and adult fish abundance over 50 years under (**A**) Scenario I (fish only), (**B**) Scenario II (fish + seals), (**C**) Scenario III (fish + seals + fishery), and (**D**) Scenario IV (full system including bold, depredating seals). Lines show median abundance across 100 demographic realisations and shaded areas show the corresponding 95% simulation intervals. Fish abundance is displayed on a logarithmic scale.

Seal persistence was comparatively insensitive to the addition of fishing or depredation. Median final seal abundance was approximately 15 individuals (9–28) in Scenario II, 15 (6– 29) in Scenario III, and 15 (4–34) in Scenario IV. Median minimum abundance was approximately five seals in Scenarios II and III, but 12 in Scenario IV. However, the underlying dynamics differed strongly. Seal populations in Scenarios II and III underwent repeated fluctuations following an initial decline (Figure 3AB), whereas in Scenario IV (Figure 3C), the non-bold adult population declined rapidly as bold adults emerged, with bold seals subsequently comprising a substantial fraction of the adult population. Thus, behavioural depredation substantially altered seal population structure without reducing overall persistence.

**Figure 3.**
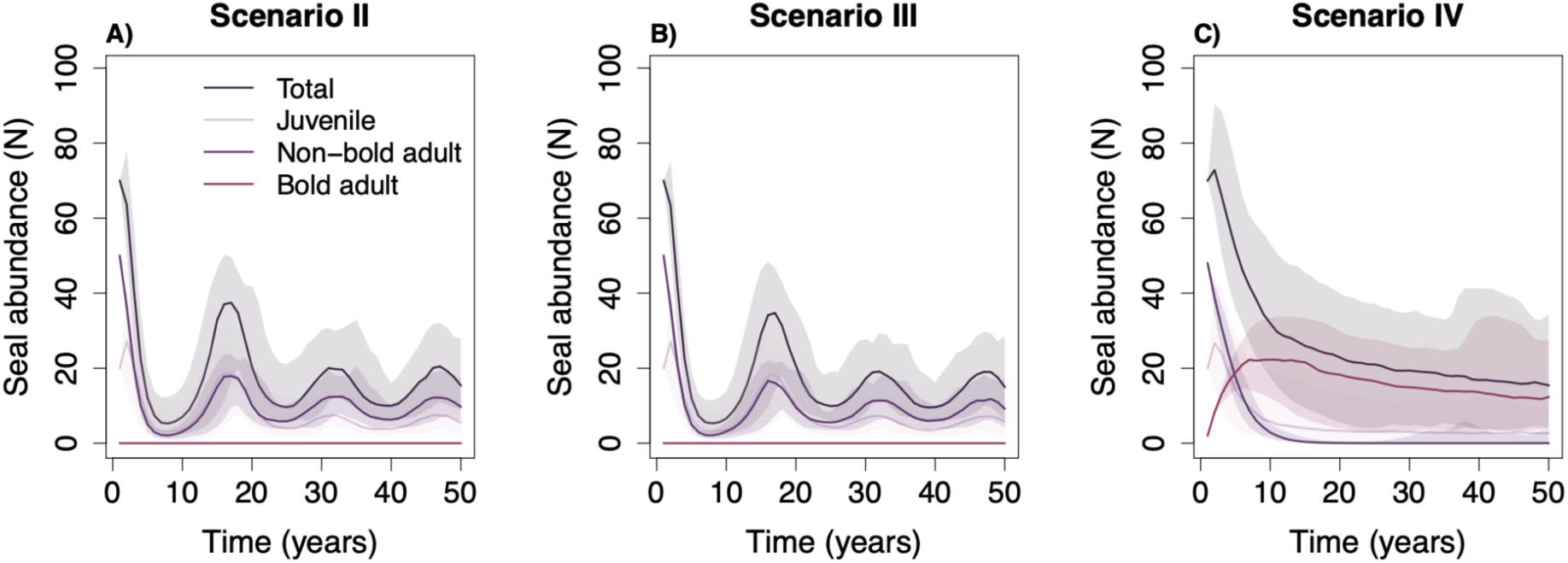
Grey seal population dynamics across scenarios with seals. Temporal dynamics of total, juvenile, non-bold adult, and bold adult grey seal abundance over 50 years under (**A**) Scenario II (fish + seals), (**B**) Scenario III (fish + seals + fishery), and (**C**) Scenario IV (full system including bold, depredating seals). Lines show median abundance across 100 demographic realisations and shaded areas show the corresponding 95% simulation intervals. Bold seals occur only in Scenario IV, in which adult seals can transition to the bold behavioural state and depredate fishery catch.

### Hypothesis 2: Non-linear fishery profitability hypothesis

We did not find support for H2, that depredation would reduce fishery profitability as increasing catch promoted the emergence of bold seals and reduced realised landings. To test H2, we compared gross catch, depredation, realised landings, and cumulative net present value (NPV) between the otherwise equivalent fishery systems without (Scenario III) and with bold seals (Scenario IV) across the 100 model realisations. Fishery production was substantially greater in the full system, despite losses to depredation (Figure 4). Median cumulative gross catch increased from 199,035 fish in Scenario III (14,719–895,930) to 875,950 in Scenario IV (11,915–2.19 million). Scenario III experienced no depredation, whereas median cumulative depredation in Scenario IV was 8.7% of gross catch (74,386 fish, 10,895–123,945). Nevertheless, median realised landings were 765,571 fish in Scenario IV compared with 199,035 in Scenario III. Bold seals reached a median final abundance of 12 individuals (3–27) in Scenario IV.

**Figure 4.**
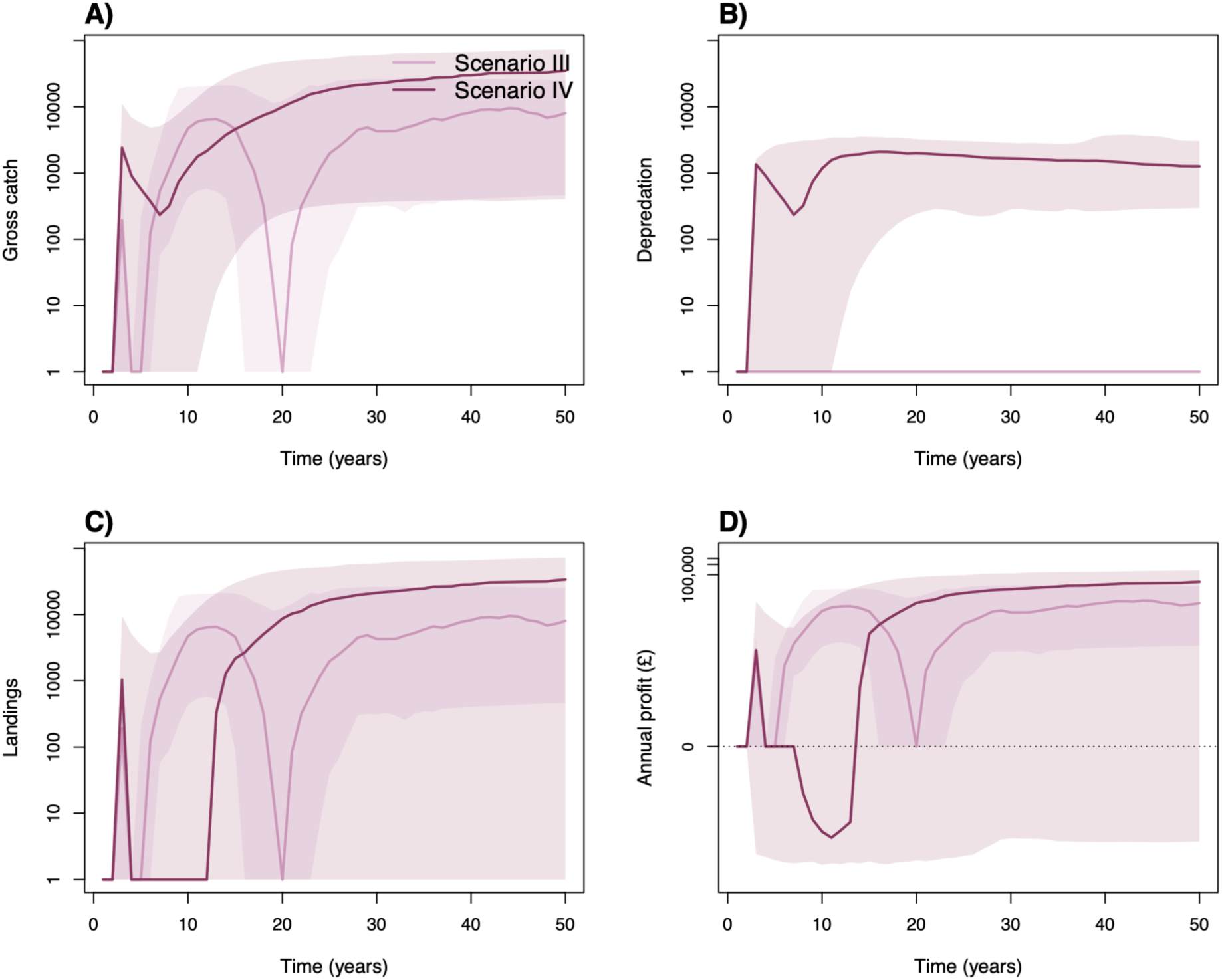
Fishery outcomes in the absence and presence of depredation. Temporal dynamics of (**A**) gross fishery catch, (**B**) catch removed through depredation, (**C**) realised landings, and (**D**) annual fishery profit over 50 years in Scenario III (fish + seals + fishery, without bold seals) and Scenario IV (full system including bold, depredating seals). Gross catch represents the number of fish harvested before depredation, whereas realised landings are the fish remaining available for sale after depredation. Lines show medians across 100 demographic realisations and shaded areas show the corresponding 95% simulation intervals. Gross catch, depredation, and landings are displayed on logarithmic scales; annual profit (£) is displayed using a signed-log transformation to accommodate the large range of positive and negative economic outcomes. Depredation is zero in Scenario III by definition.

The temporal dynamics differed strongly between scenarios (Figure 4). Scenario III showed an early increase in catch followed by a pronounced decline and subsequent recovery, whereas Scenario IV experienced an initial pulse and decline followed by a sustained increase in gross catch and landings. Depredation increased rapidly following the emergence of bold seals and subsequently remained substantial throughout the projection. These differences translated into greater long-term economic returns in the full system (Figure 5). Median cumulative NPV was £89,154 (£7,949–£435,721) in Scenario III compared with £258,799 (£1,734–£862,878) in Scenario IV. The paired comparison gave a median Scenario IV minus Scenario III difference of £116,898, although responses varied among demographic realisations: NPV was significantly greater in Scenario IV in 76 of 100 paired realisations (*t*_99_ = 9.408, *P* < 0.001), and the 95% interval of paired differences spanned -£94,245 to £511,300. Similarly, median paired realised landings were 493,177 fish greater in Scenario IV, with landings greater in 89 of 100 paired realisations. Thus, although depredation imposed a direct loss of harvested fish, this cost was generally outweighed by the greater harvesting opportunities generated by the ecological dynamics of the full system. Cumulative NPV consequently overtook Scenario III during the second half of the projection (Figure 5).

**Figure 5.**
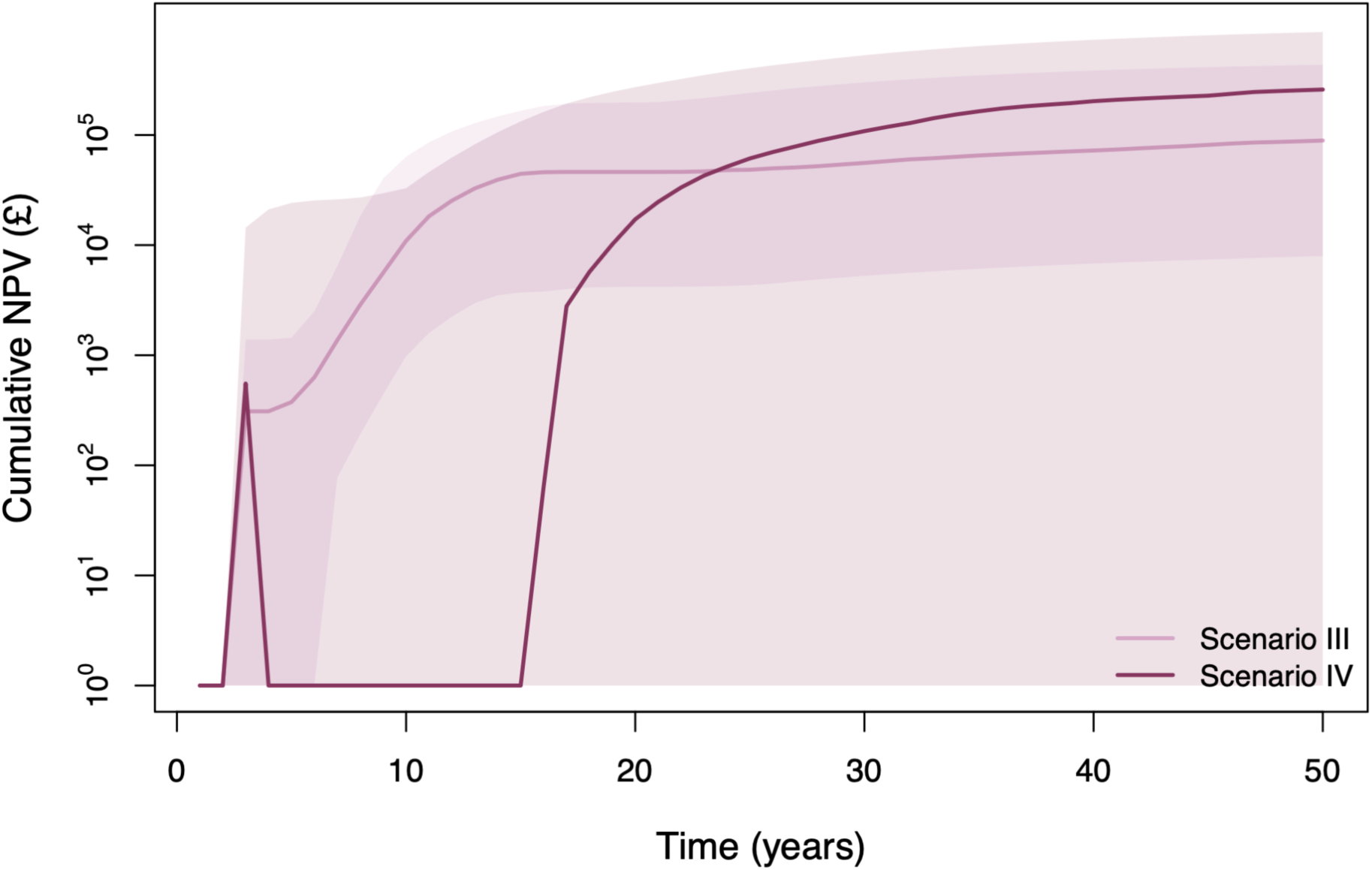
Cumulative economic performance of the fishery in the absence and presence of depredation. Cumulative net present value (NPV) of fishery profits over the 50-year projection under Scenario III (fish + seals + fishery, without bold seals) and Scenario IV (full system including bold, depredating seals). Annual profits were discounted at 5% yr^-1^ before accumulation. Lines show median cumulative NPV across 100 demographic realisations and shaded areas show the corresponding 95% simulation intervals. Cumulative NPV is displayed on a logarithmic scale to accommodate the large variation among demographic realisations.

### Hypothesis 3: System stability hypothesis

We did not find support for H3, which predicted that the emergence of bold seals would destabilise the coupled system and increase temporal variability. Instead, the full system (Scenario IV; Figure 6B) was more temporally stable than the equivalent system without depredation (Scenario III). We tested H3 by comparing coefficients of variation (CV) and variation in annual log population growth across scenarios for cod and seals. Cod temporal variability was greatest in the predator–prey system without fishing (Scenario II; Figure 6A). Median cod CV increased from 0.44 (0.24–1.20) in Scenario I to 1.00 (0.60–1.44) in Scenario II, but declined sharply after fishing was introduced, to 0.17 (0.07–0.86) in Scenario III and only 0.043 (0.039–0.046) in Scenario IV (Figure 6A). Variation in annual log population growth showed the same pattern: median SD increased from 0.17 in Scenario I to 0.41 in Scenario II, before declining to 0.038 in Scenario III and 0.0005 in Scenario IV. Thus, rather than destabilising cod dynamics, the full system produced the lowest temporal variability of all four scenarios. This pattern is also apparent in the Scenario IV population trajectory (Figure 2), which approached comparatively stable juvenile and adult abundances after its initial transient dynamics.

**Figure 6.**
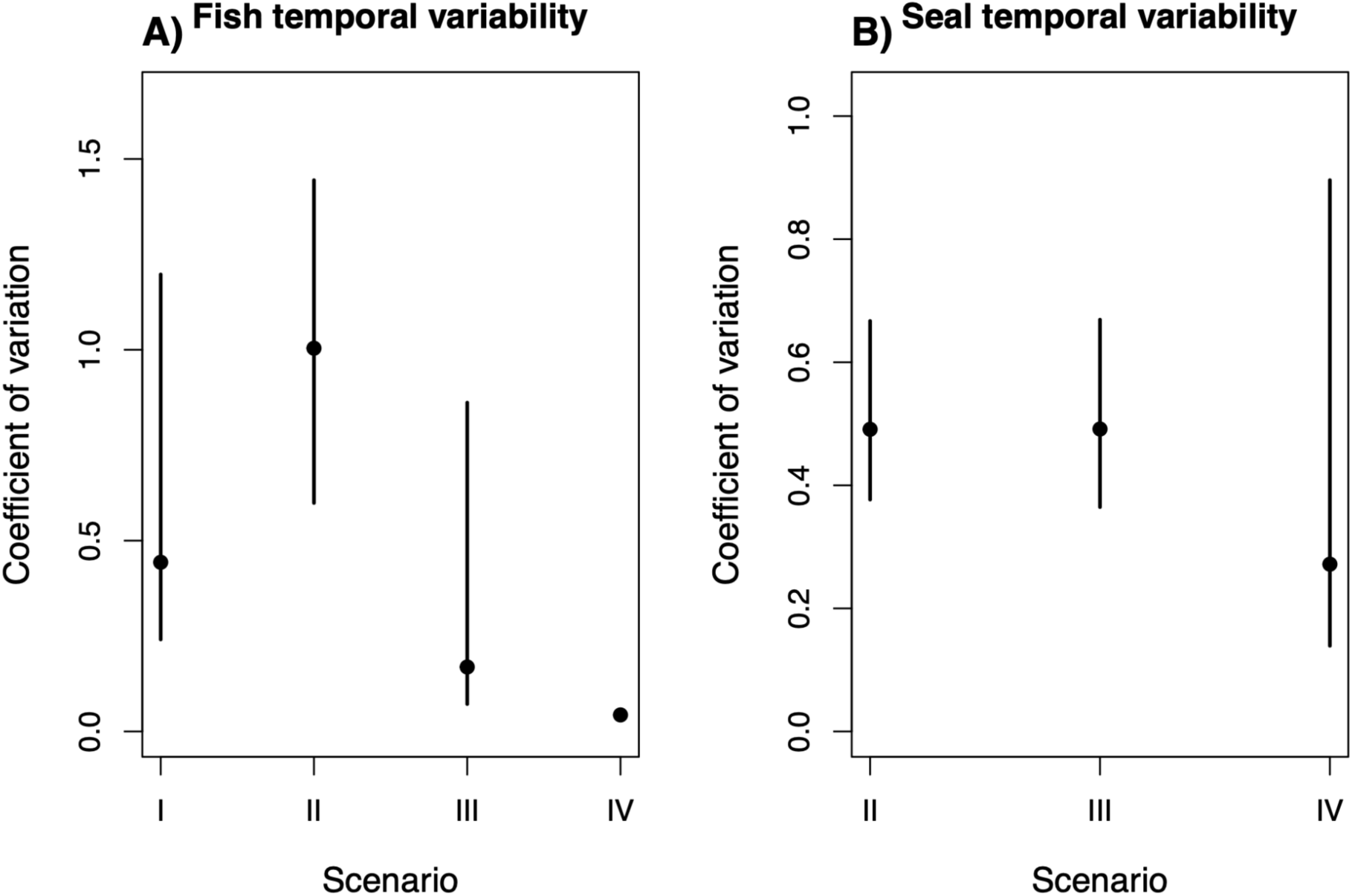
Temporal variability in fish and grey seal populations across model scenarios. Coefficients of variation in (**A**) total fish abundance and (**B**) total grey seal abundance over the 50-year projections. Fish variability is shown for Scenarios I–IV, whereas seal variability is shown for Scenarios II–IV because seals are absent from Scenario I. Points show the median coefficient of variation across 100 demographic realisations and vertical lines show the corresponding 95% simulation intervals. Larger coefficients of variation indicate greater temporal variability in population abundance.

Seal dynamics showed a similar response. Median seal CV was nearly identical in Scenarios II and III, at 0.491 (0.377–0.667 and 0.365–0.669, respectively), but declined to 0.272 (0.139–0.896) in Scenario IV ( Figure 6B). Median SD in annual log population growth similarly declined from 0.193 and 0.182 in Scenarios II and III to 0.026 in Scenario IV. Collectively, these results indicate that behavioural depredation altered the composition of the seal population and fishery outcomes but, contrary to H3, was associated with dampened rather than amplified temporal population fluctuations.

### Sensitivity analyses

Our model outcomes differed in their sensitivity to the three key assumptions we examined: bold seal energetic requirements, harvest initiation threshold, and social amplification of bold behaviour. First, increasing the proportion of bold-seal energetic requirements obtained through depredation (*m_D_*) from 0.1 to 1.0 increased median cumulative depredation by 565%, from 16,900 to 112,000 fish, while median final bold-seal abundance changed by only -1.4% (Figure S5). Median cumulative NPV declined by 45%, from £1.81 million to £0.99 million. Thus, the energetic contribution of depredated fish strongly affected the magnitude of depredation and economic returns, but had little influence on final bold seal abundance. By contrast, model outcomes were comparatively insensitive to the threshold in adult-fish growth used to initiate fishing. Increasing this threshold from 0.95 (*i.e.*, fishing occurred if the adult fish population decreased by less than 5 % relative to the adult population remaining after harvest the previous year) to 1.05 (*i.e.*, fishing occurred if the fish population increased by 5%) changed Scenario IV median cumulative NPV from £1.81 million to £1.65 million and median fish CV only from 1.53 to 1.60 (Figure S6). The trajectories across the tested thresholds were correspondingly similar for cumulative NPV, final fish abundance, temporal variability, and cumulative catch. Social amplification had a stronger effect on the behavioural composition of the seal population. Increasing the sensitivity of the non-bold-to-bold transition to existing bold seals from 0.00 to 0.40 increased median final bold-seal abundance from approximately 2 to 7 individuals and cumulative depredation from 12,100 to 20,600 fish (Figure S7). This increase in bold seals coincided with a decline in final total seal abundance as social amplification increased. Together, our sensitivity analyses indicate that the energetic value of depredated catch primarily affected economic outcomes and the quantity of fish depredated, whereas social amplification more strongly determined how extensively bold behaviour spread through the seal population.

## Discussion

Human–wildlife conflict over shared resources is intensifying as recovering wildlife populations increasingly overlap with human activities (Dirzo *et al*. 2014; Venter *et al*. 2016). Yet, such conflicts are commonly evaluated as pairwise competition in which wildlife consumption represents an equivalent loss to people (Davis *et al*. 2021; Read 2008; Tixier *et al*. 2021). This perspective overlooks a major knowledge gap: individuals differ in their propensity to exploit shared resources (Parsons *et al*. 2022; Sih *et al*. 2004; Sloan Wilson *et al*. 1994). We know comparatively little about how such behavioural heterogeneity feeds back through predator and prey demography to affect resource users. We addressed this gap by coupling stage-structured cod and grey seal demography (Caswell 2001) with harvesting, fishery economics, and transitions of individual seals into a depredating behavioural state (Holma *et al*. 2014; Punt & Donovan 2007). Three key findings stand out. First, cod and seals persisted across all scenarios despite predation and harvesting. Second, although depredation removed commercially valuable catch, the full system produced greater median long-term fishery returns than the otherwise equivalent system without depredation. Third, adding bold seals substantially stabilised rather than destabilised both cod and seal dynamics. Together, these insights show that behavioural responses to shared resources can create indirect feedbacks whose system-level consequences differ fundamentally from those expected by considering predator consumption alone.

The persistence of both species despite predation and harvesting is consistent with a large literature showing that density dependence can buffer populations against exploitation through compensatory changes in survival, reproduction, and reproduction (Brook & Bradshaw 2006; Gaillard & Yoccoz 2003; Hilborn 2013). Persistence, nevertheless, concealed substantial reorganisation of the system. Cod remained much less abundant after fishing was introduced, while allowing depredation transformed the behavioural composition of the seal population without appreciably changing its final total abundance. This distinction is important because human–wildlife interactions are increasingly recognised as being driven disproportionately by subsets of individuals rather than homogeneous populations (Graham *et al*. 2011b; Sih *et al*. 2004). Fisheries provide particularly strong opportunities for such specialisation because gear concentrates prey into predictable and accessible patches, potentially rewarding repeated interaction with humans (Jackson *et al*. 2024). Evidence from killer whales further demonstrates that depredation can develop through both individual and social learning (Auguin *et al*. 2025; Earl *et al*. 2021). Thus, predator abundance alone may be a poor proxy to quantify *factual* conflict intensity: two predator populations of similar size can interact very differently with people if their behavioural composition differs.

The economic consequences of our study similarly challenge the commonly held assumption that the cost of depredation is equivalent to the market value of the fish removed (Tanner & Davis 2026). Depredation generated a genuine direct loss in our model because harvesting costs were incurred before seals removed fish. Yet, this action did not translate into lower long-term median profitability. Such counterintuitive outcomes are familiar from multispecies bioeconomic theory, where changes in predator abundance, prey abundance, and harvesting can propagate through food webs in non-linear ways so that immediate and equilibrium economic effects differ in sign (Clark 1993) (Ryan *et al*. 2010). Ecosystem-based fishery models likewise show that maximising catch, economic rent, and predator–prey persistence need not identify the same system state or management strategy (Costalago *et al*. 2019). Our simulations extend this reasoning by making predator behaviour itself endogenous to fishing: harvesting not only removes cod but creates feeding opportunities that change seal behaviour, which subsequently alters predator and prey demography and future harvesting opportunities. Consequently, the economic effect of depredation is better understood as a system-level counterfactual than as the price of individual fish lost from nets. This integrative perspective does not imply that depredation generally benefits fisheries, not least because a quarter of our paired realisations generated lower returns in the full system, but it does show why direct-loss calculations alone can misrepresent its longer-term consequences.

The more complex scenario in our models was the most stable. Although ecological complexity was classically expected to destabilise communities (May 1972), subsequent theory and experiments have shown that additional species (Loreau & de Mazancourt 2013), states (Giménez Romero *et al*. 2025), and interactions (McCann *et al*. 1998) can instead dampen fluctuations when they create compensatory responses, asynchronous dynamics or appropriately distributed interaction strengths (Lehman & Tilman 2000; McCann 2000; Tilman *et al*. 2006). This result also fits naturally within Chesson’s coexistence framework (Chesson 2000). Stabilising mechanisms promote coexistence when ecological differences cause populations to experience stronger effective limitations when common than when rare, while temporal environmental variation can additionally promote coexistence through mechanisms such as the storage effect and relative nonlinearity (Barabás *et al*. 2018; Chesson 2003). In our model, introducing a bold behavioural state increases the number of demographic pathways through which density dependence, food availability, and harvesting feed back into the system. Importantly, juvenile seal performance depends on the abundance of seals across behavioural states, not simply non-bold adults, while bold and non-bold adults experience partly distinct resource- and density-dependent feedbacks. We therefore suggest that partitioning adults between behavioural states distributes rather than simply adds interaction pressure, generating compensatory feedbacks capable of damping the predator–prey oscillations apparent in the simpler system. We note that this is a mechanistic hypothesis emerging from the model rather than a formal decomposition of Chesson’s coexistence mechanisms, but it provides a clear explanation for why increasing behavioural complexity need not reduce stability.

There are many precedents for greater complexity producing greater stability. In experimental grasslands, increasing plant diversity increased temporal stability of ecosystem production because fluctuations among species partly compensated for one another, producing portfolio and overyielding effects (Lehman & Tilman 2000; Tilman *et al*. 2006). In food webs, weak and intermediate trophic interactions can dampen consumer–resource oscillations and reduce the probability that populations approach extinction (Huxel & McCann 1998; McCann *et al*. 1998). Similarly, modest external energetic inputs can stabilise trophic dynamics when consumers continue to rely primarily on endogenous resources, although sufficiently large subsidies can reverse this effect (Newsome *et al*. 2015; Oro *et al*. 2013). The general principle is not that complexity is intrinsically stabilising, but that adding partially asynchronous components creates opportunities for one component to compensate when another declines, the basis of ecological insurance theory (Yachi & Loreau 1999). Bold and non-bold seals can be interpreted similarly as alternative demographic–behavioural pathways through which the predator population responds to changing natural and anthropogenic resources. The marked reduction in variability in our full model may therefore represent an intraspecific analogue of the stabilising effects of functional diversity observed at the community level.

Our findings have important implications for human perceptions of depredating predators. Fisheries conflicts are rarely experienced as abstract system-level dynamics: fish removed from a net are immediate, visible losses (Davis *et al*. 2021; Tanner & Davis 2026), making the individual responsible an obvious focus for frustration and management action (Davis *et al*. 2021; Read 2008; Tixier *et al*. 2021). Bold predators (here, bold seals) may consequently be perceived by people (here, fishers) as particularly damaging individuals because their costs are conspicuous whereas any stabilising ecological effects are diffuse, delayed and difficult to observe, let alone quantify. Similar asymmetries between perceived and system-level impacts occur widely in human–wildlife conflict, where salient losses caused by particular animals can dominate attitudes even when population-level ecological effects are more complicated (Treves *et al*. 2016). Our findings do not negate fishers’ experience of depredation: a fish removed from gear remains an immediate, real economic loss. Instead, our findings caution against translating said immediate loss into the conclusion that depredating individuals are necessarily detrimental to the overall system and to the fishery. In our model, the very behavioural state that imposed those visible costs formed part of a more stable system that maintained greater cod abundance and, in most realisations, greater long-term economic returns. Managing bold individuals solely as “problem animals” could therefore remove interactions that have indirect stabilising consequences that are invisible at the scale of an individual fishing event.

Our study emphasises the need for a different approach to managing complex human-wildlife conflict. Attempts to resolve depredation often focus on reducing predator abundance (Davis *et al*. 2021), removing problem individuals (Graham *et al*. 2011b), or deterring predators from fishing gear (Guerra 2019), despite mixed evidence for the long-term effectiveness of such interventions (Barrios-Guzmán *et al*. 2024). If system behaviour depends on behavioural composition and feedback structure rather than predator abundance alone, removing bold seals need not produce proportional ecological or economic benefits. Conversely, interventions that modify the opportunity structure generating depredation, such as gear modifications, reduced soak times or changes in the spatial and temporal predictability of fishing, may weaken the energetic reward and behavioural reinforcement without removing predators (Auguin *et al*. 2025; Jackson *et al*. 2024). The sensitivity of our model to social amplification reinforces this point: behavioural transmission affected how extensively boldness spread, whereas the energetic contribution of depredation more strongly affected economic losses. Management should therefore distinguish between reducing the number of depredating individuals, reducing the incentive to depredate, and reducing the economic consequences when depredation occurs. These are mechanistically different objectives and need not have equivalent ecological outcomes.

Several assumptions limit how literally these dynamics should be interpreted. We represented depredation as a discrete and irreversible behavioural state, although empirical studies demonstrate substantial individual and temporal heterogeneity in interactions with fisheries (Earl *et al*. 2021; Graham *et al*. 2011b). We also excluded additional mortality experienced by depredating animals through entanglement, bycatch, or deliberate killing, despite evidence that interaction with fisheries can simultaneously provide energetic opportunities and increase mortality risk (Davis 2022; Magera *et al*. 2013). Finally, future expansions of our modelling framework should explicitly explore the role of prey switching, spatial structure, and adaptive redistribution of fishing effort on predator–prey and economic dynamics. These expansions are particularly relevant to the apparent stabilising mechanism: our interpretation that behavioural partitioning creates compensatory demographic pathways should be tested explicitly by decomposing the feedbacks among juvenile, non-bold, and bold adult seals and by perturbing the dependence of juvenile vital rates on total seal abundance. Spatial extensions incorporating haul-out sites, cod distributions, and adaptive fisher movement (van Putten *et al*. 2012) would provide a further test of whether stabilisation persists when behavioural states also differ spatially.

More broadly, our findings suggest that behavioural diversity deserves consideration alongside species and functional diversity when evaluating the stability of human-modified ecosystems. The ecological insurance theory predicts that aggregate dynamics can become more stable when different components respond asynchronously to environmental change (Yachi & Loreau 1999), while coexistence theory emphasises that stabilising differences in ecological responses can allow interacting types to persist despite competition (Chesson 2000). Human activities increasingly create precisely these new behavioural niches, from crop raiding and livestock depredation to scavenging on waste and exploitation of fishing gear (Hoare 1999; Newsome *et al*. 2015; Treves *et al*. 2016). Treating the individuals that exploit those niches simply as competitors or “problem animals” risks overlooking their position within a larger network of compensatory ecological feedbacks. For fisheries, coexistence with recovering marine predators may therefore depend not only on how many fish each actor removes, but on how fishing itself creates behavioural opportunities and how those behaviours reshape the stability and economic performance of the coupled system.

## Supporting information

SOM

## Acknowledgements

This publication arises from research funded by the John Felel Oxford University Press Research Fund (0012020) to K.J.D.. R.S-G. was supported by NERC Pushing the Frontiers grant (NE/X013766/1). We thank the contributors of seal and fish demographic data in the COMADRE Animal Matrix Database.

## Author contributions

K.J.D. conceived the study, designed the bioeconomic framework, and parameterised the fisheries components; R.S-G. and K.J.D. co-designed and implemented the demographic modelling and phylogenetic imputations; both authors contributed to the interpretation of results. K.J.D. led the first draft of the manuscript, and R.S-G. provided revisions. Both authors gave final approval for publication.

## Data availability statement

All data and commented R scripts used in this study will be openly available on Zenodo upon publication. Demographic data for fish and seals were obtained from the open-access COMADRE Animal Matrix Database (v.4.21.1; https://www.comadre-db.org). Body mass data for fishes were sourced from FishBase (https://www.fishbase.org).

