## Supplementary material for "Wildlife depredation stabilizes ecological-economic systems and can lead to improved economic outcomes": SOM

12 **Table S1.** Source of demographic information used of the 65 fish species used to impute the vital rates of the Atlantic cod (*Gadus morhua*).
13 Vital rates obtained from the COMADRE Animal Matrix Database (Salguero-Gomez *et al.* 2016). Vital rates are: $\sigma_{f_j}$: juvenile survival, $\gamma_{f_j}$:
14 maturation, $\sigma_{f_a}$: adult survival, and $\phi_{f_a}$: adult reproduction. Year: year of publication. Body mass (in grams) obtained from FishBase
15 (Boettiger et al. 2012).

| Latin name | Common name | Family | Authors | Journal | DOI/ISBN | Year | $\sigma_{f_j}$ | $\gamma_{f_j}$ | $\sigma_{f_a}$ | $\phi_{f_a}$ | Body mass |
| --- | --- | --- | --- | --- | --- | --- | --- | --- | --- | --- | --- |
| *Acipenser fulvescens* | Lake sturgeon | Acipenseridae | Vélez-Espino; Koops | N Am J Fish Manage | 10.1577/M08-034.1 | 2009 | 0.027 | 1.000 | 0.295 | 65.545 | 125000.00 |
| *Acipenser transmontanus* | White sturgeon | Acipenseridae | Vélez-Espino; Koops | Ecol Model | 10.1016/j.ecolmodel.2012.09.022 | 2012 | 0.000 | 1.000 | 0.766 | 16804.847 | 816000.00 |
| *Ambloplites rupestris* | Rock bass | Centrarchidae | Peoples | Master Thesis | NA | 2010 | 0.246 | 1.000 | 0.060 | 24.295 | 1360.00 |
| *Ammocrypta pellucida* | Eastern sand darter | Percidae | Vélez-Espino; Koops | Ecol Model | 10.1016/j.ecolmodel.2012.09.022 | 2012 | 0.020 | 1.000 | 0.262 | 37.204 | 2.50 |
| *Astroblepus ubidiai* | Andean catfish | Astroblepidae | Vélez-Espino | Ecol Freshw Fish | 10.1111/j.1600-0633.2005.00084.x | 2005 | 0.175 | 1.000 | 0.419 | 5.306 | 10.00 |
| *Brachyrhaphis rhabdophora* | Live-bearing fish | Poeciliidae | Johnson; Zúñiga-Vega | Ecology | 10.1890/07-1672.1 | 2009 | 0.810 | 0.432 | 0.833 | 0.718 | 1.20 |
| *Catostomus catostomus* | Salish sucker | Catostomidae | Vélez-Espino; Koops | Ecol Model | 10.1016/j.ecolmodel.2012.09.022 | 2012 | 0.000 | 1.000 | 0.297 | 1512.211 | 3300.00 |
| *Catostomus platyrhynchus* | Mountain sucker | Catostomidae | Young; Koops | Fish & Oceans Can | NA | 2013 | 0.004 | 1.000 | 0.240 | 104.861 | 500.00 |
| *Chasmistes liorus* | NA | Catostomidae | Tuckfield; Belk | Popul Ecol | 10.1002/1438-390X.12005 | 2019 | 0.001 | 1.000 | 0.888 | 11.966 | 2500.00 |
| *Chrosomus oreas* | Mountain redbelly dace | Cyprinidae | Peoples | Master Thesis | NA | 2010 | 0.064 | 1.000 | 0.004 | 718.000 | 6.40 |
| *Clinocottus analis* | Woolly sculpin | Cottidae | Davis; Levin | Mar Ecol Prog Ser | 10.3354/meps234229 | 2002 | 0.860 | 0.849 | 0.837 | 0.317 | 80.00 |
| *Clinostomus elongatus* | Redside dace | Cyprinidae | Vélez-Espino; Koops | Ecol Model | 10.1016/j.ecolmodel.2012.09.022 | 2012 | 0.004 | 1.000 | 0.263 | 202.123 | 10.00 |
| *Clinostomus funduloides* | Rosyside dace | Cyprinidae | Peoples | Master Thesis | NA | 2010 | 0.135 | 1.000 | 0.018 | 1098.000 | 10.00 |
| *Coregonus huntsmani* | Atlantic whitefish | Salmonidae | Vélez-Espino; Koops | Ecol Model | 10.1016/j.ecolmodel.2012.09.022 | 2012 | 0.000 | 1.000 | 0.627 | 6402.675 | 1000.00 |
| *Coregonus reighardi* | Shortnose cisco | Salmonidae | Vélez-Espino; Koops | Ecol Model | 10.1016/j.ecolmodel.2012.09.022 | 2012 | 0.000 | 1.000 | 0.590 | 1466.864 | 539.00 |
| *Coregonus zenithicus* | Shortjaw cisco | Salmonidae | Vélez-Espino; Koops | Ecol Model | 10.1016/j.ecolmodel.2012.09.022 | 2012 | 0.000 | 1.000 | 0.620 | 1832.257 | 300.00 |
| *Cottus bairdii* | Mottled sculpin | Cottidae | Peoples | Master Thesis | NA | 2010 | 0.524 | 1.000 | 0.266 | 30.655 | 125.00 |
| *Cottus confusus* | Shorthorn sculpin | Cottidae | Vélez-Espino; Koops | Ecol Model | 10.1016/j.ecolmodel.2012.09.022 | 2012 | 0.007 | 1.000 | 0.469 | 72.037 | 45.00 |
| *Cottus duranii* | Eastslope sculpin | Cottidae | Vélez-Espino; Koops | Ecol Model | 10.1016/j.ecolmodel.2012.09.022 | 2012 | 0.010 | 1.000 | 0.440 | 55.046 | 40.00 |
| *Cyprinus carpio* | Common carp | Cyprinidae | Stratford; Pollino; Brown | Environ Modell Softw | 10.1016/j.envsoft.2016.02.009 | 2016 | 0.500 | 1.000 | 0.650 | 2.381 | 40090.00 |
| *Epinephelus morio* | Red grouper | Serranidae | Fujiwara; Zhou | Can J Fish Aquat Sci | 10.1139/cjfas-2012-0520 | 2013 | 0.869 | 0.059 | 0.869 | 0.140 | 23000.00 |
| *Erimyzon sucetta* | Lake chub sucker | Catostomidae | Vélez-Espino; Koops | Ecol Model | 10.1016/j.ecolmodel.2012.09.022 | 2012 | 0.000 | 1.000 | 0.830 | 2706.999 | 400.00 |
| *Etheostoma flabellare* | Fantail darter | Percidae | Peoples | Master Thesis | NA | 2010 | 0.313 | 1.000 | 0.097 | 139.458 | 4.00 |
| *Gasterosteus aculeatus* | Enos lake stickleback | Gasterosteidae | Vélez-Espino; Koops | Ecol Model | 10.1016/j.ecolmodel.2012.09.022 | 2012 | 0.008 | 1.000 | 0.094 | 11.985 | 1.50 |
| *Genypterus blacodes* | Pink cusk-eel | Ophidiidae | Gonzales-Olivares; Aranguiz-Acuna; Ramos-Jiliberto; Rojas-Palma | Fish Res | 10.1016/j.fishres.2008.11.006 | 2009 | 0.989 | 1.000 | 0.795 | 0.311 | 25000.00 |
| *Hybognathus argyritus* | Western Silvery Minnow | Cyprinidae | Vélez-Espino; Koops | Ecol Model | 10.1016/j.ecolmodel.2012.09.022 | 2012 | 0.002 | 1.000 | 0.184 | 334.222 | 100.00 |
| *Hypseleotris klunzingeri* | Carp gudgeon | Eleotridae | Yen; Bond; Shenton; Spring; MacNally | J Appl Ecol | 10.1111/1365-2664.12074 | 2013 | 0.150 | 1.000 | 0.100 | 100.000 | 5.00 |
| *Lepisosteus oculatus* | Spotted gar | Lepisosteidae | Vélez-Espino; Koops | Ecol Model | 10.1016/j.ecolmodel.2012.09.022 | 2012 | 0.000 | 1.000 | 0.743 | 5240.616 | 4440.00 |
| *Maccullochella peelii* | Murray cod | Percichthyidae | Stratford; Pollino; Brown | Environ Modell Softw | 10.1016/j.envsoft.2016.02.009 | 2016 | 0.466 | 1.000 | 0.723 | 1.378 | 113500.00 |
| *Macquaria ambigua* | Golden perch | Percichthyidae | Yen; Bond; Shenton; Spring; MacNally | J Appl Ecol | 10.1111/1365-2664.12074 | 2013 | 0.530 | 1.000 | 0.576 | 13.791 | 24000.00 |
| *Macrhybopsis storeriana* | Carmine shiner | Cyprinidae | Young; Koops | Fish & Oceans Can | NA | 2013 | 0.000 | 1.000 | 0.393 | 1391.914 | 150.00 |
| *Micropterus dolomieu* | Smallmouth bass | Centrarchidae | Spromberg; Birge | Environ Toxicol Chem | 10.1897/04-160.1 | 2005 | 0.040 | 1.000 | 0.307 | 20.951 | 5410.00 |
| *Morone saxatilis* | Striped bass | Moronidae | Vélez-Espino; Koops | Ecol Model | 10.1016/j.ecolmodel.2012.09.022 | 2012 | 0.000 | 1.000 | 0.684 | 34119.321 | 57000.00 |
| *Moxostoma duquesnii* | Black redhorse | Catostomidae | Vélez-Espino; Koops | Ecol Model | 10.1016/j.ecolmodel.2012.09.022 | 2012 | 0.000 | 1.000 | 0.703 | 1297.294 | 2000.00 |
| *Moxostoma hubbsi* | Copper redhorse | Catostomidae | Vélez-Espino; Koops | Ecol Model | 10.1016/j.ecolmodel.2012.09.022 | 2012 | 0.000 | 1.000 | 0.799 | 11253.474 | 1800.00 |
| *Neogobius melanostomus* | Round goby | Gobiidae | Spromberg; Birge | Environ Toxicol Chem | 10.1897/04-160.1 | 2005 | 0.030 | 1.000 | 0.108 | 33.570 | 381.42 |
| *Nocomis leptcephalus* | Bluehead chub | Cyprinidae | Peoples | Master Thesis | NA | 2010 | 0.122 | 1.000 | 0.015 | 388.918 | 50.00 |
| *Notropis anogenus* | Pugnose shiner | Cyprinidae | Vélez-Espino; Koops | Ecol Model | 10.1016/j.ecolmodel.2012.09.022 | 2012 | 0.007 | 1.000 | 0.141 | 126.900 | 3.00 |
| *Notropis percobromus* | Carmine shiner | Cyprinidae | Vélez-Espino; Koops | Ecol Model | 10.1016/j.ecolmodel.2012.09.022 | 2012 | 0.008 | 1.000 | 0.235 | 95.998 | 4.00 |
| *Notropis photogenis* | Silver shiner | Cyprinidae | Young; Koops | Fish & Oceans Can | NA | 2012 | 0.001 | 1.000 | 0.119 | 151.204 | 5.00 |
| *Noturus stigmosus* | Northern madtom | Ictaluridae | Vélez-Espino; Koops | Ecol Model | 10.1016/j.ecolmodel.2012.09.022 | 2012 | 0.011 | 1.000 | 0.372 | 55.800 | 60.00 |
| *Oncorhynchus clarkii lewisi* | Westslope cutthroat trout | Salmonidae | Vélez-Espino; Koops | Ecol Model | 10.1016/j.ecolmodel.2012.09.022 | 2012 | 0.008 | 1.000 | 0.664 | 39.930 | 2500.00 |
| *Oncorhynchus gilae* | Gila trout | Salmonidae | Brown; Echelle; Propst; | West N Am Naturalist | NA | 2001 | 0.491 | 1.000 | 0.149 | 8.633 | 1200.00 |
|  |  |  | Brooks;<br>Fisher |  |  |  |  |  |  |  |  |
| *Oncorhynchus kisutch* | Coho salmon | Salmonidae | Spromberg; Birge | Environ Toxicol Chem | 10.1897/04-160.1 | 2005 | 0.050 | 1.000 | 0.069 | 20.022 | 15170.00 |
| *Oncorhynchus tshawytscha* | Chinook salmon | Salmonidae | Wilson | Conserv Biol | 10.1046/j.1523-1739.2003.01535.x | 2003 | 0.059 | 1.000 | 0.621 | 4.480 | 61400.00 |
| *Opsopoeodus emiliae* | Pugnose minnow | Cyprinidae | Young; Koops | Fish & Oceans Can | - | 2012 | 0.010 | 1.000 | 0.202 | 14.458 | 8.00 |
| *Osmerus spectrum* | Utopia dwarf smelt | Osmeridae | Vélez-Espino; Koops | Ecol Model | 10.1016/j.ecolmodel.2012.09.022 | 2012 | 0.001 | 1.000 | 0.348 | 4380.620 | 50.00 |
| *Percina copelandi* | Channel darter | Percidae | Vélez-Espino; Koops | Ecol Model | 10.1016/j.ecolmodel.2012.09.022 | 2012 | 0.005 | 1.000 | 0.214 | 164.352 | 10.00 |
| *Pimephales promelas* | Fathead minnow | Cyprinidae | Gleason | Hum Ecol Risk Assess | 10.1080/20018091094835 | 2001 | 0.001 | 1.000 | 0.390 | 1121.110 | 2.00 |
| *Poecilia reticulata* | Trinidadian guppy | Poeciliidae | Bronikowski; Clark; Rodd; Reznick | Ecology | 10.1890/0012-9658(2002)083[2194:PDCOPI]2.0.CO;2 | 2002 | 0.670 | 0.179 | 0.720 | 1.144 | 0.25 |
| *Pterois miles* | Common lionfish | Scorpaenidae | Morris; Shertzer; Rice | Biol Invasions | 10.1007/s10530-010-9786-8 | 2011 | 0.000 | 1.000 | 0.876 | 9776.131 | 1000.00 |
| *Pterois volitans* | Red lionfish | Scorpaenidae | Morris; Shertzer; Rice | Biol Invasions | 10.1007/s10530-010-9786-8 | 2011 | 0.000 | 1.000 | 0.876 | 9776.131 | 1442.00 |
| *Pyloodictis olivaris* | Flathead catfish | Ictaluridae | Sakaris; Irwin | Ecol Appl | 10.1890/08-0305.1 | 2010 | 0.000 | 1.000 | 0.848 | 2769.130 | 55790.00 |
| *Retropinna semoni* | Australian smelt | Retropinnidae | Yen; Bond; Shenton; Spring; MacNally | J Appl Ecol | 10.1111/1365-2664.12074 | 2013 | 0.300 | 1.000 | 0.050 | 8.000 | 3.00 |
| *Rhinichthys cataractae* | Nooksack dace | Cyprinidae | Vélez-Espino; Koops | Ecol Model | 10.1016/j.ecolmodel.2012.09.022 | 2012 | 0.001 | 1.000 | 0.531 | 335.287 | 20.00 |
| *Rhinichthys osculus* | Speckled dace | Cyprinidae | Vélez-Espino; Koops | Ecol Model | 10.1016/j.ecolmodel.2012.09.022 | 2012 | 0.006 | 1.000 | 0.208 | 123.844 | 15.00 |
| *Rutilus rutilus* | Common roach | Cyprinidae | Otjacques; De Laender; Kestemont | Ecol Model | 10.1016/j.ecolmodel.2015.12.002 | 2016 | 0.019 | 1.000 | 0.489 | 26.322 | 1840.00 |
| *Salvelinus confluentus* | Bull trout | Salmonidae | Bowerman | - | - | 2013 | 0.218 | 1.000 | 0.306 | 2.479 | 14510.00 |
| *Salvelinus fontinalis* | Brook char | Salmonidae | Vélez-Espino; Koops | Ecol Model | 10.1016/j.ecolmodel.2012.09.022 | 2012 | 0.001 | 1.000 | 0.456 | 546.698 | 8000.00 |
| *Salvelinus malma* | Lake trout | Salmonidae | Spromberg; Birge | Environ Toxicol Chem | 10.1897/04-160.1 | 2005 | 0.050 | 1.000 | 0.245 | 18.278 | 18300.00 |
| *Sardina pilchardus* | Sardine | Clupeidae | Serghini; Boutayeb; Auger; Charouki; Ramzi; | Acta Biotheo | 10.1007/s10441-009-9090-0 | 2009 | 0.520 | 0.519 | 0.190 | 3.060 | 150.00 |
|  |  |  | Ettahiri;<br>Tchente |  |  |  |  |  |  |  |  |
|  |  |  | Haslob;<br>Hauss;<br>Petereit; |  |  |  |  |  |  |  |  |
| *Sprattus sprattus* | European sprat | Clupeidae | Clemmesen;<br>Kraus; Peck | Mar Biol | 10.1007/s00227-012-1933-6 | 2012 | 0.937 | 0.057 | 0.866 | 0.293 | 8.50 |
| *Zingel asper* | Percid | Percidae | Labonne;<br>Gaudin | Can J Fish<br>Aquat Sci | 10.1139/f05-245 | 2006 | 0.500 | 1.000 | 0.500 | 1.500 | 100.00 |
|  |  |  | Bergek; Ma;<br>Vetemaa;<br>Franzén;<br>Appelberg | Ecotox<br>Environ Safe | 10.1016/j.ecoenv.2012.01.019 | 2012 | 0.026 | 1.000 | 0.580 | 22.643 | 510.00 |
| *Zoarces viviparus* | European eelpout | Zoarcidae |  |  |  |  |  |  |  |  |  |
| *Gadus morhua* | Atlantic cod | Gadidae | - | - | - | - | - | - | - | - | 96000.00 |

17 **Table S2.** Estimates of phylogenetic inertia for the four vital rates and adult body mass of the
18 65 fish species used to impute vital rates of the Atlantic cod (*Gadus morhua*) in our study.

| Trait | Blomberg's K | P-value | Pagel's $\lambda$ | P-value |
| --- | --- | --- | --- | --- |
| Juvenile survival ($\sigma_{f_j}$) | 0.106 | 0.001 | 0.826 | 0.013 |
| Maturation ($\gamma_{f_j}$) | 0.235 | 0.001 | 0.984 | <0.001 |
| Adult survival ($\sigma_{f_a}$) | 0.068 | 0.029 | 0.911 | <0.001 |
| Reproduction ($\phi_{f_a}$) | 0.098 | 0.002 | 0.803 | 0.105 |
| Body mass | 0.208 | 0.001 | 0.927 | <0.001 |

19

20 **Table S3.** Mean (and 95% C.I.) of the vital rates of the examined fish species' population

| Vital rate | Mean | 5% C.I. | 95% C.I. | 21 dynamics,<br>22 Atlantic<br>23 cod<br>24 (*Gadus*<br>*25 morhua*),<br>26 after 40 |
| --- | --- | --- | --- | --- |
| Juvenile survival ($\sigma_{f_j}$) | 0.0157 | 0.001 | 0.114 | |
| Maturation ($\gamma_{f_j}$) | 0.914 | 0.788 | 1.000 | |
| Adult survival ($\sigma_{f_a}$) | 0.750 | 0.525 | 0.901 | |
| Reproduction ($\phi_{f_a}$) | 30.115 | 7.372 | 42.000 | |

27 phylogenetic imputations.

28
29
30

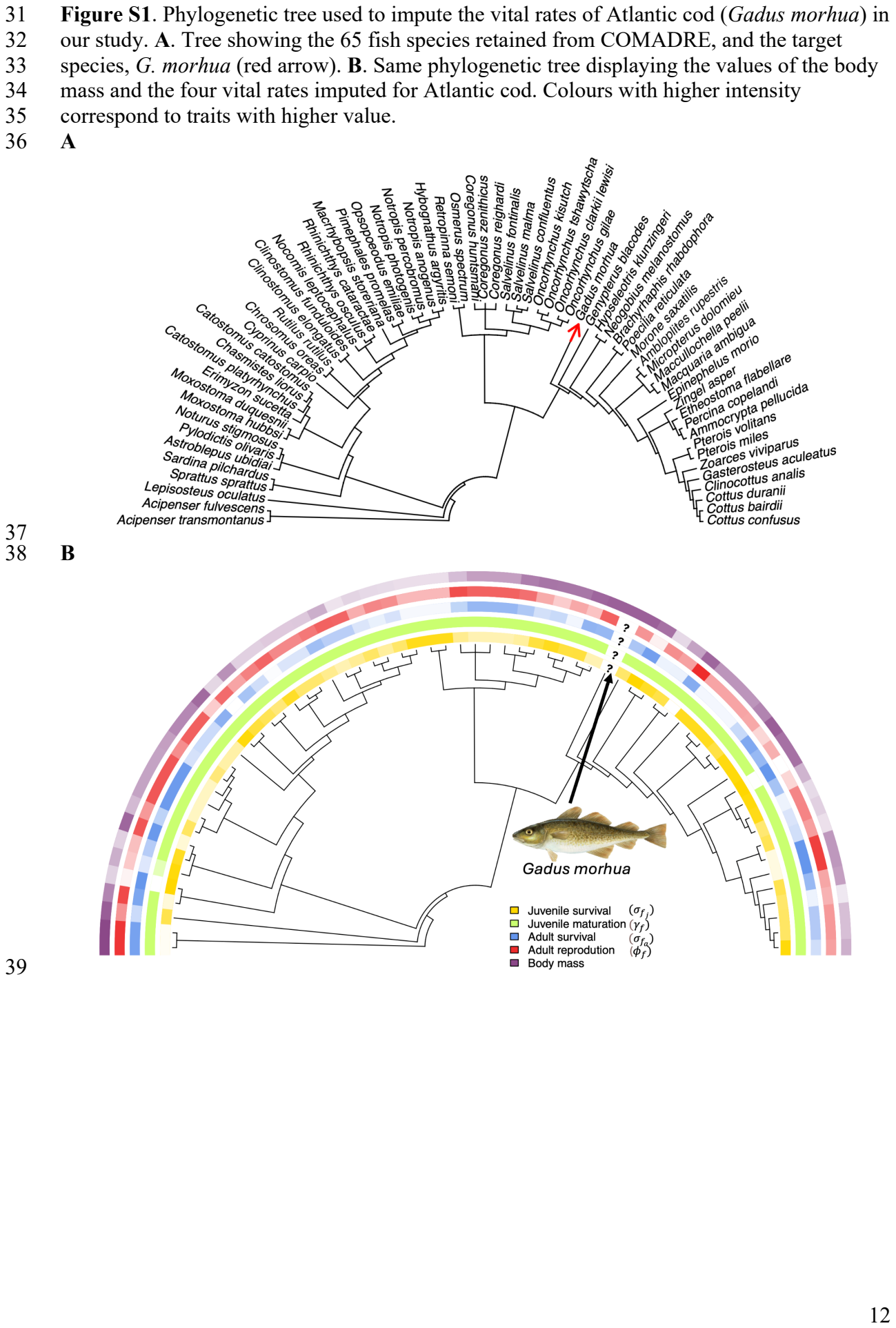

**Figure S2.** Matrix dimension of the 65 fish species collected from the COMADRE Animal Matrix Database to impute vital rates for *Gadus morhua*. Note that the majority (n = 39) were already at 2×2 dimensions, and thus did not require collapsing.

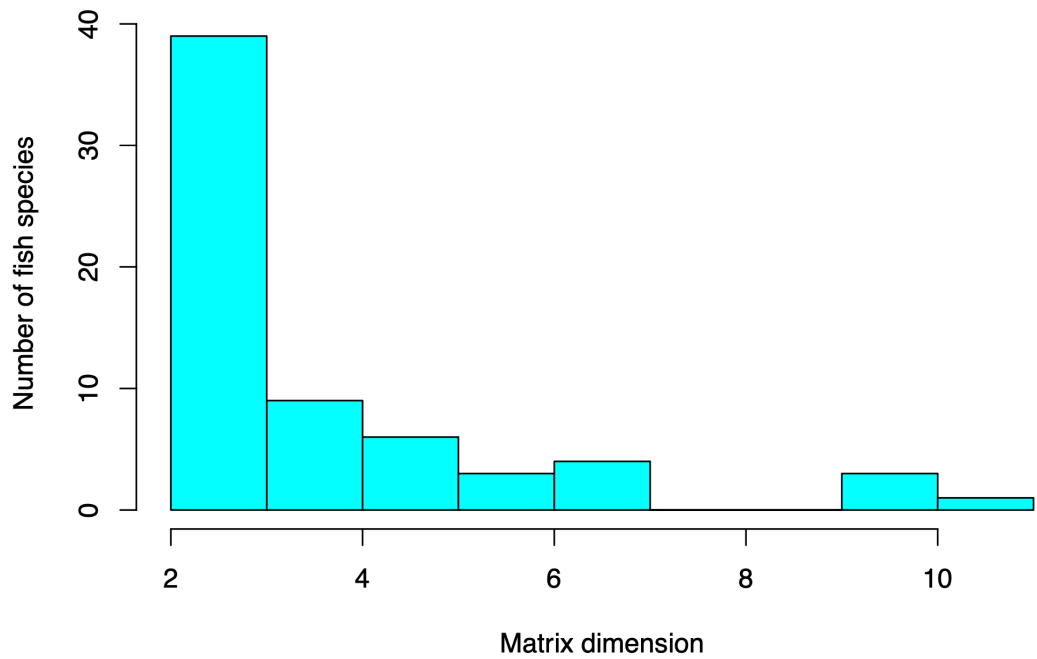

**Figure S3.** Distribution of key demographic outputs of the imputed 100 matrix population models for *Gadus morhua*.

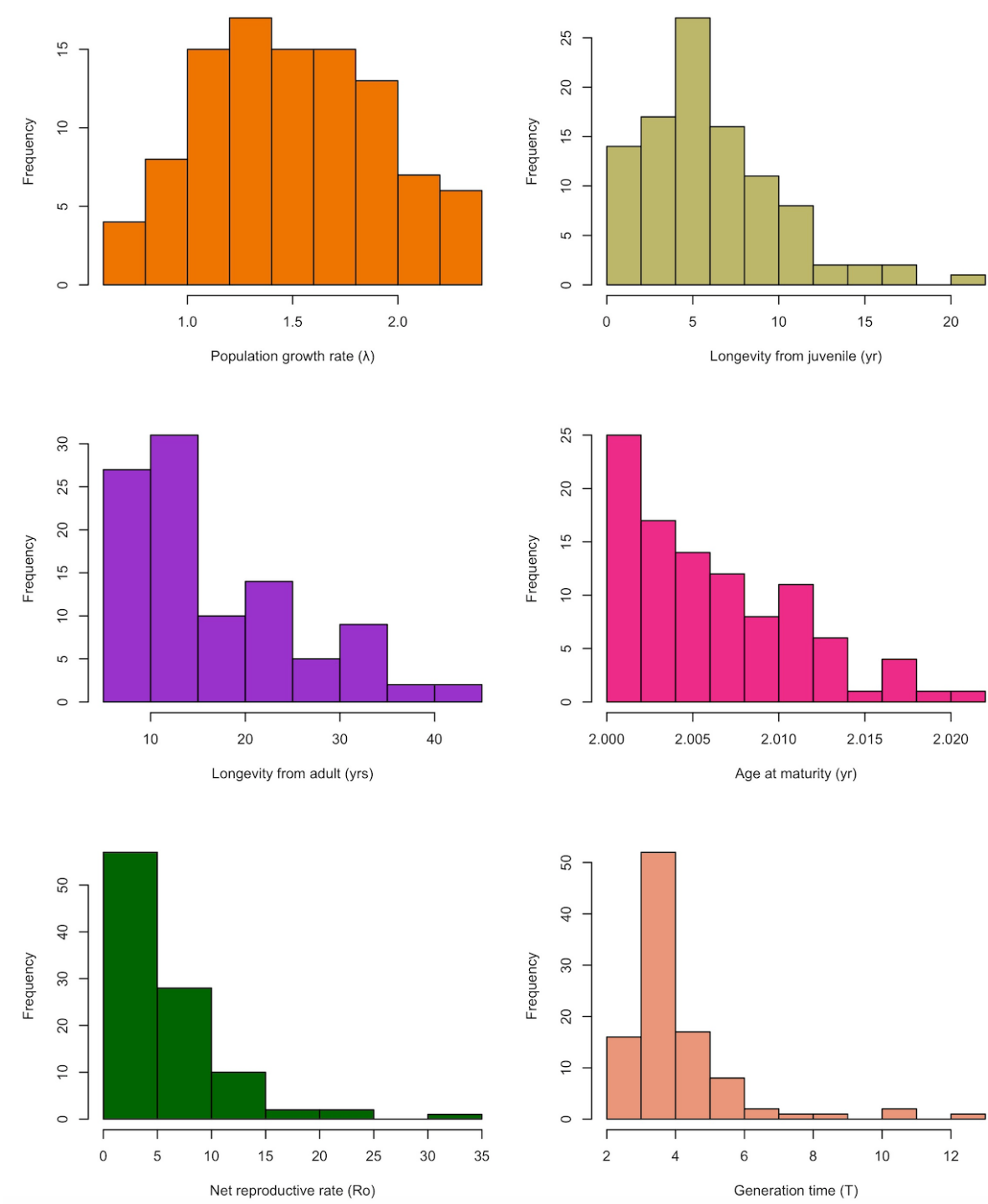

**Figure S4.** Demographic comparison of the building blocks of the matrix population model without (gray) and with (orange) adult bold seals. A) Juvenile survival, B) juvenile maturation, C) Adult survival, D) Adult reproduction, and E) Population growth rate.

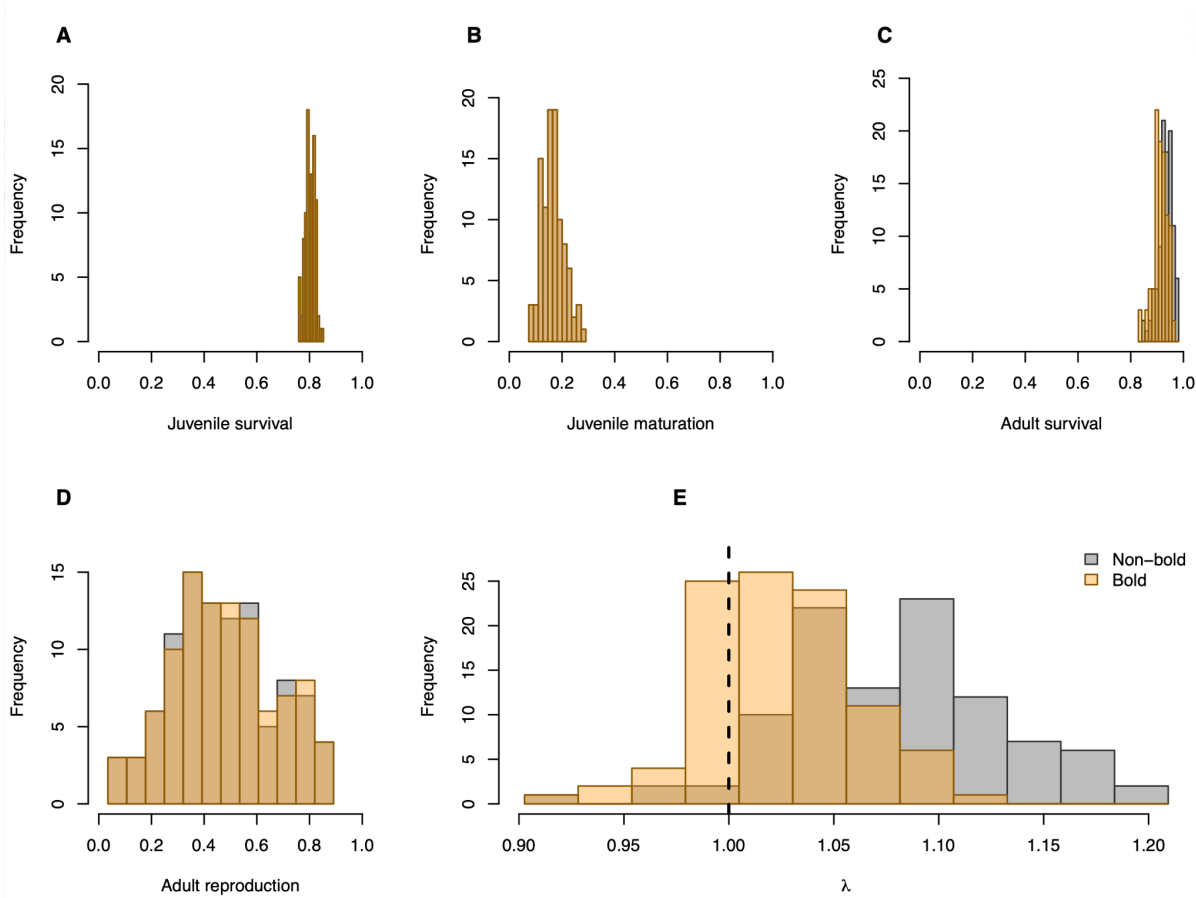

**Figure S5.** Sensitivity of ecological and economic outcomes to the energetic contribution of depredation. Sensitivity of cumulative fishery net present value (NPV), cumulative catch removed through depredation, final bold-seal abundance, and temporal variability in fish abundance to $m_D$, the proportion of a bold seal's annual energetic requirements obtained through depredation. The parameter $m_D$ was varied from 0.1 (the baseline parameterisation) to 1.0 in increments of 0.1 while all other parameters were held at their baseline values. Points and lines show medians across 100 realisations and shaded areas show the 95% C.I. Cumulative NPV and cumulative depredation are displayed on logarithmic scales. Fish temporal variability is expressed as the coefficient of variation in total fish abundance over the 50-year projection.

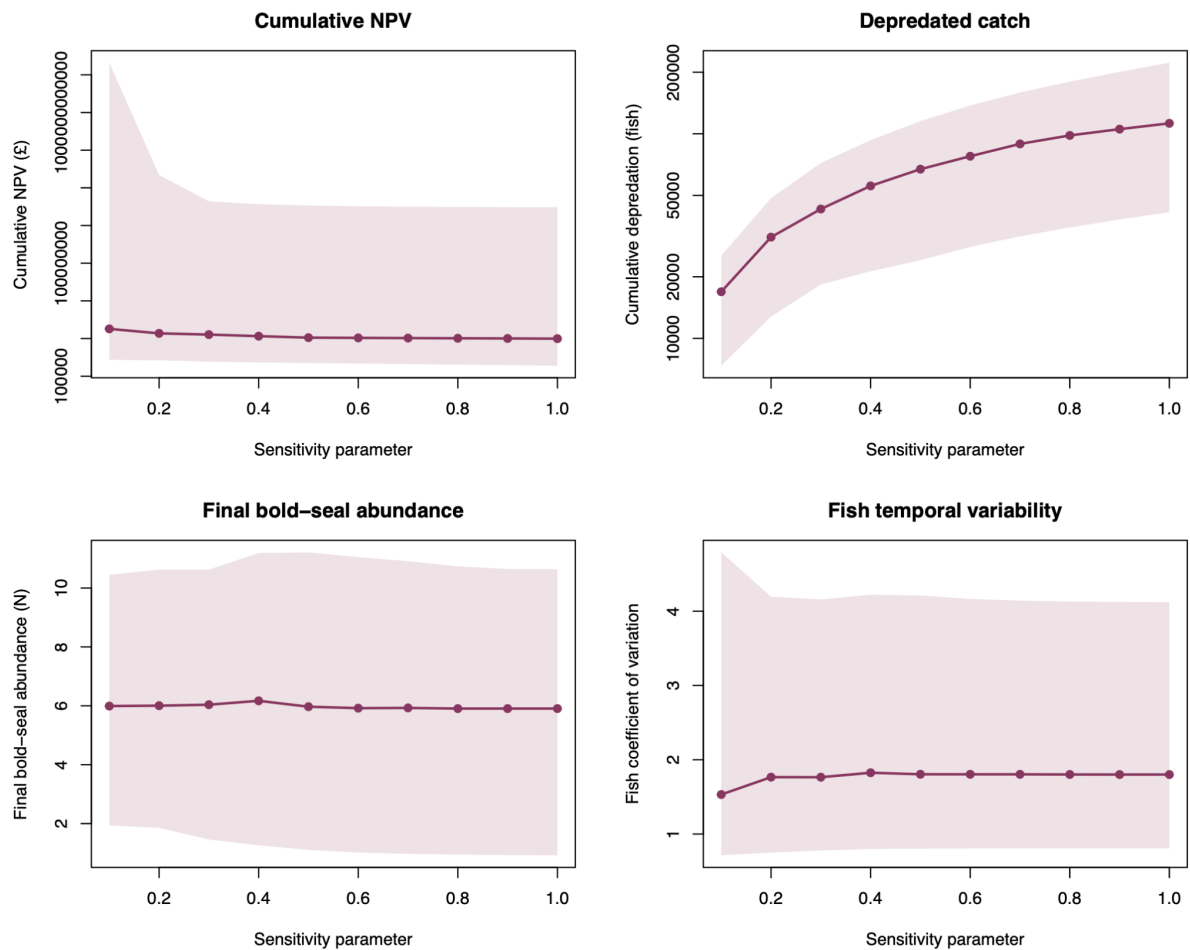

**Figure S6.** Sensitivity of ecological and economic outcomes to the harvest initiation threshold. Sensitivity of cumulative fishery net present value (NPV), final fish abundance, temporal variability in fish abundance, and cumulative fishery catch to the adult-fish population growth threshold at which harvesting was initiated. Thresholds of 0.95, 1.00 (baseline), and 1.05 represent progressively more restrictive conditions for initiating harvest. Outcomes are shown for Scenario III (fish + seals + fishery, without bold seals) and Scenario IV (full system including bold, depredating seals), where applicable. Points and lines show medians across 100 demographic realisations and shaded areas show the corresponding 95% simulation intervals. Cumulative NPV, final fish abundance, and cumulative catch are displayed on logarithmic scales; temporal variability is expressed as the coefficient of variation in total fish abundance over the 50-year projection.

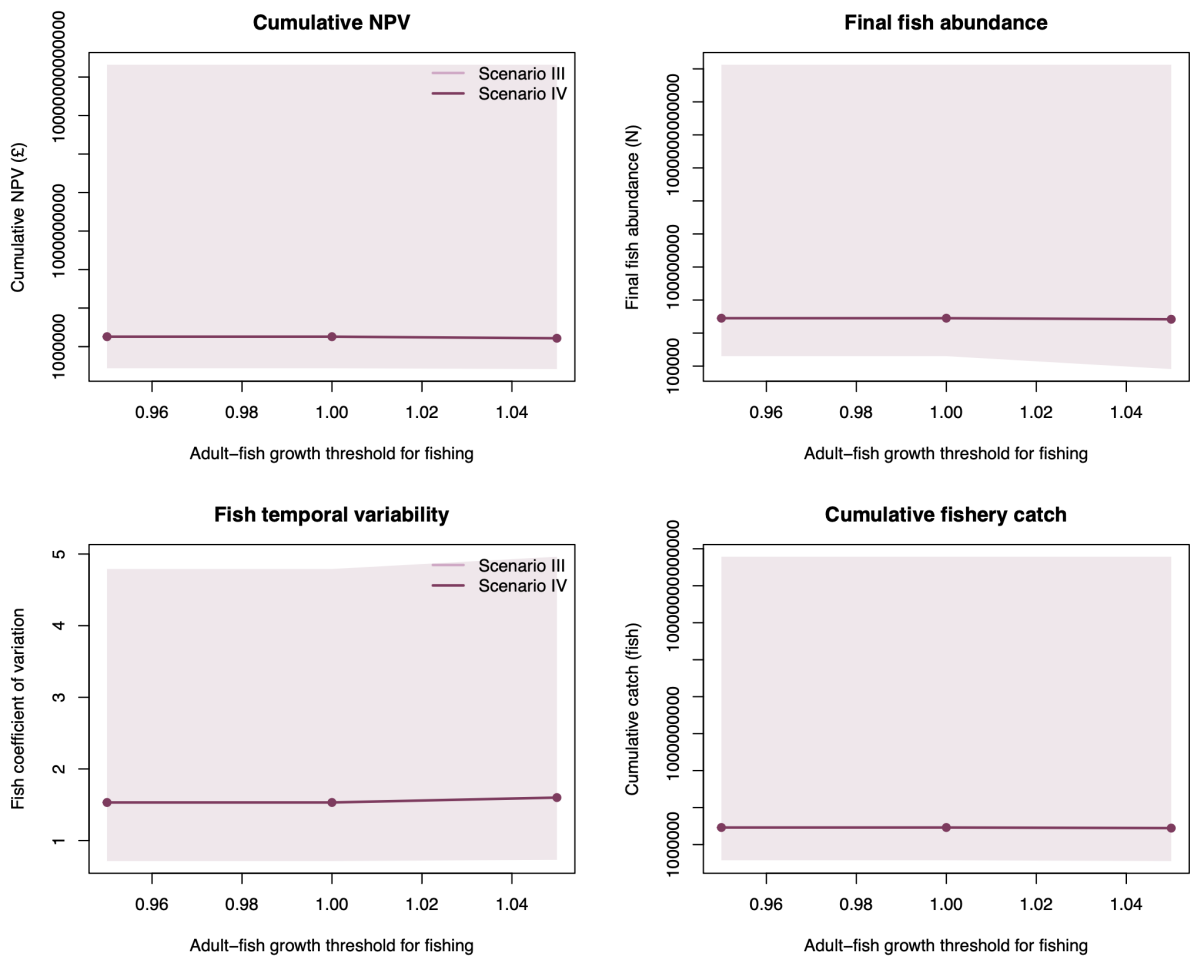

**Figure S7.** Sensitivity of ecological and economic outcomes to social amplification of depredation behaviour. Sensitivity of final bold-seal abundance, cumulative catch removed through depredation, cumulative fishery net present value (NPV), and final total seal abundance to the strength of the positive effect of existing bold seals on the transition of non-bold adults into the bold behavioural state. The sensitivity parameter was varied among 0.00 (no social or frequency-dependent amplification), 0.50 (baseline parameterisation), and 1.00 (strong amplification), while all other parameters were held at their baseline values. Points and lines show medians across 100 realisations and shaded areas show the corresponding 95% C.I. Cumulative depredation and NPV are displayed on logarithmic scales. The sensitivity represents the hypothesised social or frequency-dependent component of the transition to depredation behaviour in Eq. 32.

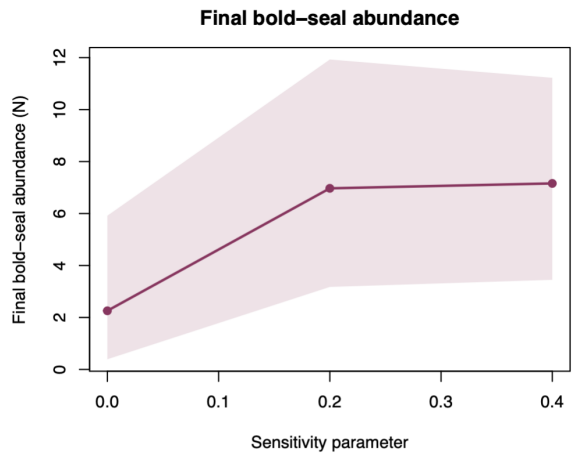

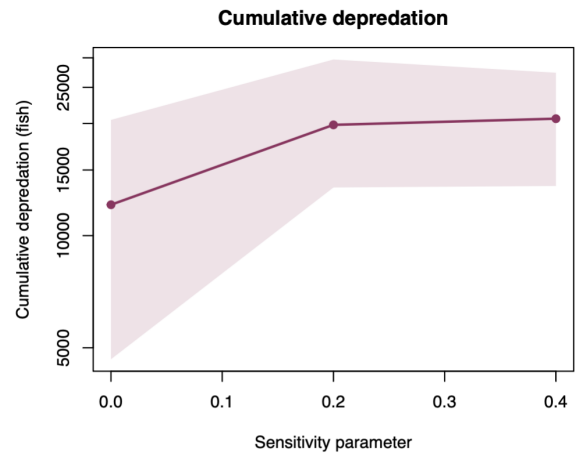

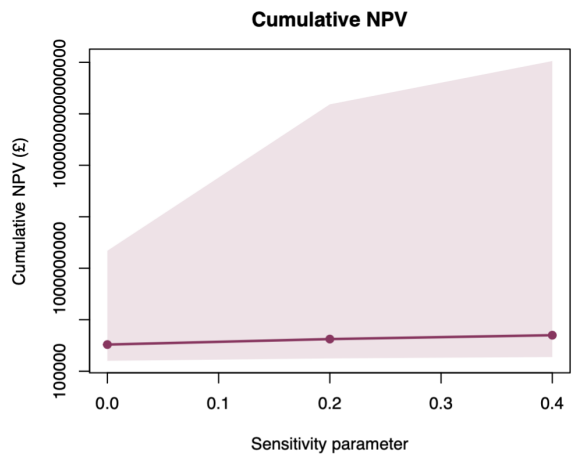

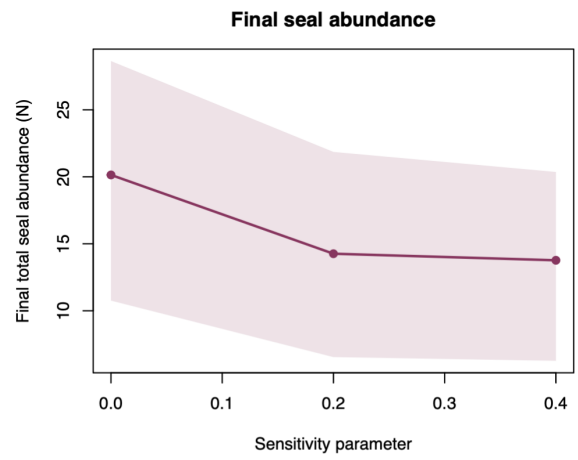
